# MetaWorm: An Integrative Data-Driven Model Simulating *C. elegans* Brain, Body and Environment Interactions

**DOI:** 10.1101/2024.02.22.581686

**Authors:** Mengdi Zhao, Ning Wang, Xinrui Jiang, Xiaoyang Ma, Haixin Ma, Gan He, Kai Du, Lei Ma, Tiejun Huang

## Abstract

The behavior of an organism is profoundly influenced by the complex interplay between its brain, body, and environment. Existing data-driven models focusing on either the brain or the body-environment separately. A model that integrates these two components is yet to be developed. Here, we present MetaWorm, an integrative data-driven model of a widely studied organism, *C. elegans*. This model consists of two sub-models: the brain model and the body & environment model. The brain model was built by multi-compartment models with realistic morphology, connectome, and neural population dynamics based on experimental data. Simultaneously, the body & environment model employed a lifelike body and a 3D physical environment, facilitating easy behavior quantification. Through the closed-loop interaction between two sub-models, MetaWorm faithfully reproduced the realistic zigzag movement towards attractors observed in *C. elegans*. Notably, MetaWorm is the first model to achieve seamless integration of detailed brain, body, and environment simulations, enabling unprecedented insights into the intricate relationships between neural structures, neural activities, and behaviors. Leveraging this model, we investigated the impact of neural system structure on both neural activities and behaviors. Consequently, MetaWorm can enhance our understanding of how the brain controls the body to interact with its surrounding environment.

## Introduction

The behaviors of an organism are not simply the product of brain activity but rather, emerge from a dynamic interplay among the brain, body, and environment. To unravel the underlying neural control mechanisms, it is crucial to develop an integrative simulation that incorporates detailed models of the brain, body and environment, to both validate theories and make predictions for biological experiments. Currently, there are specific data-driven models that simulate either the brain, e.g., the mouse primary visual cortex^1^, the mouse striatum^2^, and the rat somatosensory cortex^3^, or exclusively the body and environment to reproduce animal behaviors, such as neuromechanical models for Drosophila^4^ and rodent^5^. OpenWorm project^6^ aimed to simulate *C. elegans* years ago, despite its detailed brain model and body model worked separately. So far, no data-driven model exists that successfully integrates these elements, although such integration is urgently needed to progress our understanding of neural control mechanisms^7^.

An integrative data-driven model must meet certain requirements. First, it should incorporate a biophysically detailed brain model that has similar neural structures and neural activity patterns as real organism. Secondly, it should consist of a realistic and high-performance body and environment model that facilitates easy behavior quantification. Thirdly, and most crucially, the brain model should not only control the body model interacting with the virtual environment, but also receive the feedback from the body and environment model, establishing a closed-loop interaction^8,9^.

*C. elegans* stands out as an exemplary model for developing such an integrative data-driven model that bridges brain, body and environment. It has completely mapped morphologies of all 302 neurons^10,11^ (in adult hermaphrodite), along with the connectome and synapse-level structure^12-14^. Recordings of single neural activity^15-23^, as well as brain-wide neural dynamics^18,20,24^ are available. Its body is well-reconstructed, consisting of only 95 muscle cells. It also exhibits easily quantifiable behaviors, such as crawling, swimming and foraging^25,26^. Therefore, these simple structures and abundant data provide solid foundations for creating an integrative data-driven model.

In this study, we present an integrative data-driven model of *C. elegans*, MetaWorm (Fig. 1). First, we developed a neural network model (brain model) of a *C. elegans* foraging neural circuit. It was built using biophysically detailed compartmental neuronal models, with compartments under 2 μm in length, encoding neural activity similar to live *C. elegans*. Second, we developed a body & environment model of *C. elegans*. The body model contained 96 muscles and enabled efficient real-time simulation at 30 frames per second and easy quantification of behaviors. Third, we established a closed-loop interaction between the brain model and the body-environment model to simulate *C. elegans* moving towards an attractor in a zigzag shape. Sensory neurons in the neural network model were activated by attractor’s concentration in environment. Muscles in the body model were controlled by motor neurons in the neural network model. Finally, we performed synthetic perturbations on the neural network model and suggested that neurite or synaptic/gap junction’ absence disrupts global neural dynamics and hampers accurate forward motion. This highlights the importance of network structure in shaping both network dynamics and behavior patterns.

**Fig 1.**
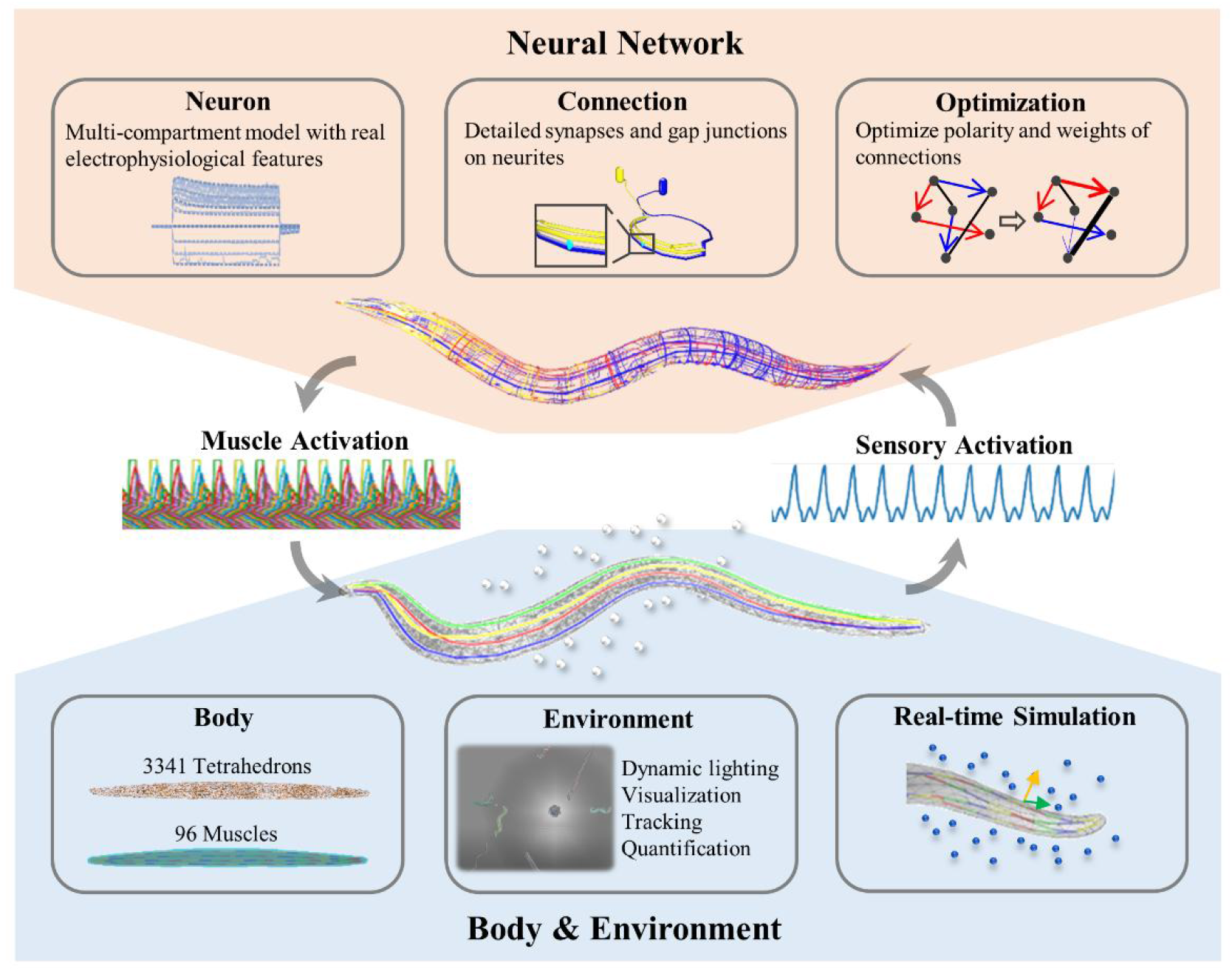
Overview of MetaWorm. MetaWorm comprises two sub-models: the neural network model and the body & environment model. The neural network model is a biophysically detailed. Its neuron models are multi-compartmental, exhibiting electrophysiological features similar to real *C. elegans* neurons. These neuron models are connected by synapses and gap junctions on neurites. An optimization tool is employed to tune the weights and polarities of these connections. The body & environment model consists of a biomechanical body, a 3D fluid environment and a real-time simulation engine. The soft body is constructed using 3341 tetrahedrons, with 96 muscles arranged from head to tail as actuators. The large-scale 3D environment allows for the integration of various agents and elements including *C. elegans*, E. coli (food), and fluid. And it provides a range of auxiliary tools for dynamic lighting, visualization, tracking, and behavior analysis. Real-time simulation is achieved through a Finite Element Method (FEM) solver and simplified hydrodynamics. To achieve foraging behavior in *C. elegans*, the neural network senses attractor’s concentration in the environment, performs neural computations to control muscles’ contraction and relaxation, and ultimately deforms the body to propel it towards the attractor.

MetaWorm is open-source and modular as the brain, body, environment models and their interactions can be independently modified, improved or extended by the scientific community. MetaWorm will contribute to a deeper understanding of the intricate relationships between neural structures, neural activities, and behaviors. This work also provides a valuable framework for developing brain-body-environment model of other organisms. This innovative approach paves the way for new avenues of systems biology and has the potential to revolutionize traditional neural mechanism research.

## Results

### Constructing a data-driven biophysically detailed neural network model of *C. elegans*

Our neural network model of *C. elegans* contained 136 neurons that participated in sensory and locomotion functions, as indicated by published studies^24,27-31^. To construct this model, we first collected the necessary data including neural morphology, ion channel models, electrophysiology of single neurons, connectome, connection models and network activities (Fig. 2A). Next, we constructed the individual neuron models and their connections (Fig. 2B). Subsequently, a biophysically detailed model without functions was established (Fig. 2C). Finally, we optimized the weights and polarities of the connections to obtain a model that reflected realistic dynamics (Fig. 2D). An overview of the model construction is illustrated in Figure 2.

**Fig 2.**
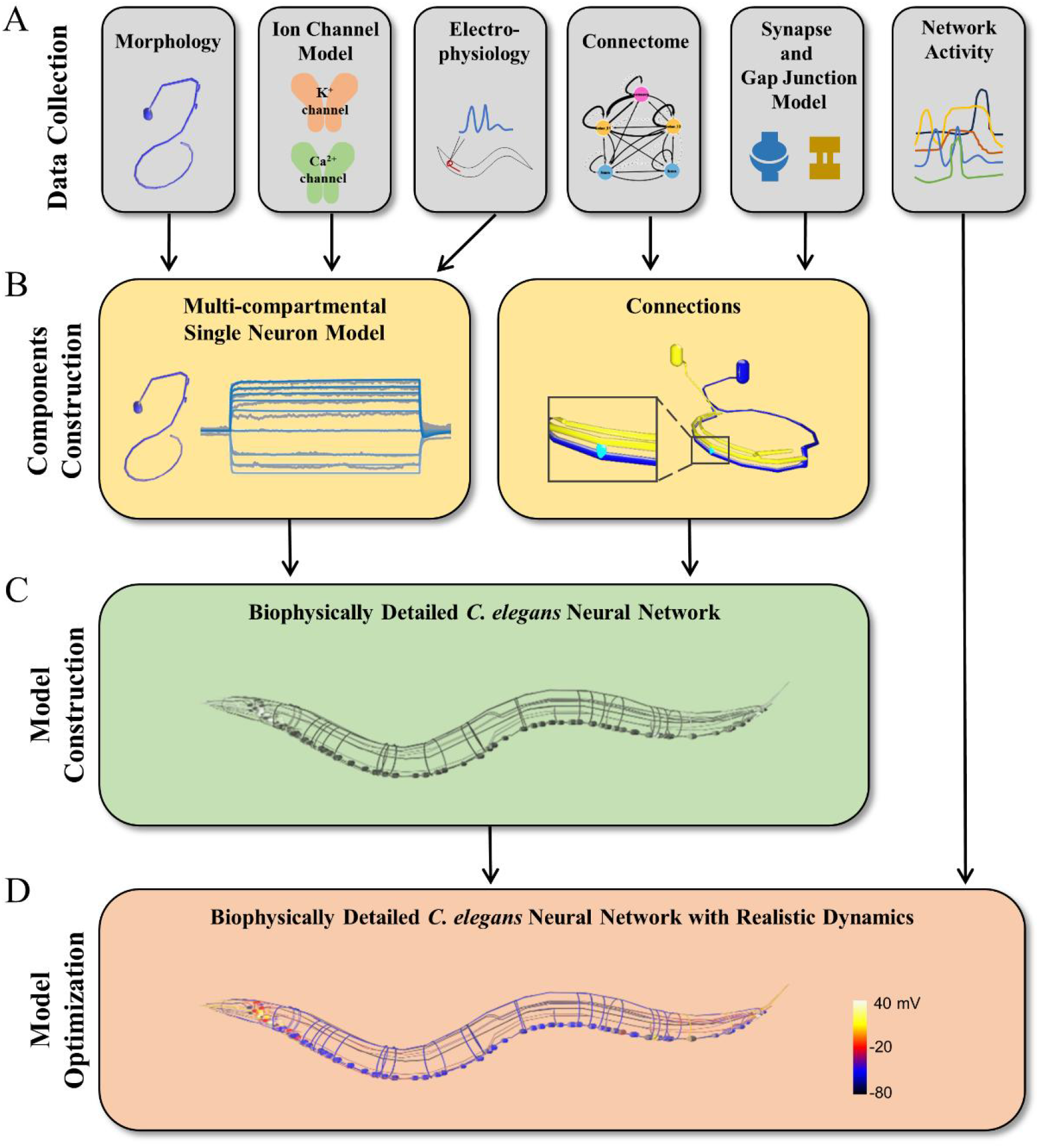
Construction of biophysically-detailed neural network model of *C. elegans*. **A**. Experimental data collection to constrain models. These data include neural morphologies, ion channel models, electrophysiological recordings of single neurons, connectome, connection models, and neural network activities. **B**. Construction of multi-compartmental neuron models and connection models (synapses and gap junctions). **C**. The biophysically detailed *C. elegans* neural network model without functional neural activities. **D**. Optimization of the biophysically detailed *C. elegans* neural network model to achieve realistic network dynamics. Neurons are color-coded to represent membrane potential.

To achieve a high level of biophysical and morphological realism in our model, we employed multi-compartment models to represent individual neurons. The morphologies of neuron models were constructed based on published morphological data^32,33^. Soma and neurite sections were further divided into several segments, where each segment was less than 2 μm in length. We combined 14 known classes of ion channels (Supplementary Table S1-2)^34^ in neuron models, and tuned the passive parameters and ion channel conductance densities by an optimization algorithm^35^ to reproduced electrophysiological traces in the experiments^36-39^. Based on the few available electrophysiological data, we digitally reconstructed models of five representative neurons: AWC, AIY, AVA, RIM and VD5. The I-V curves of our neuron models perfectly match that in the experiments (Fig. 3). The optimized parameters of neuron models are listed in Supplementary Table S3.

**Fig 3.**
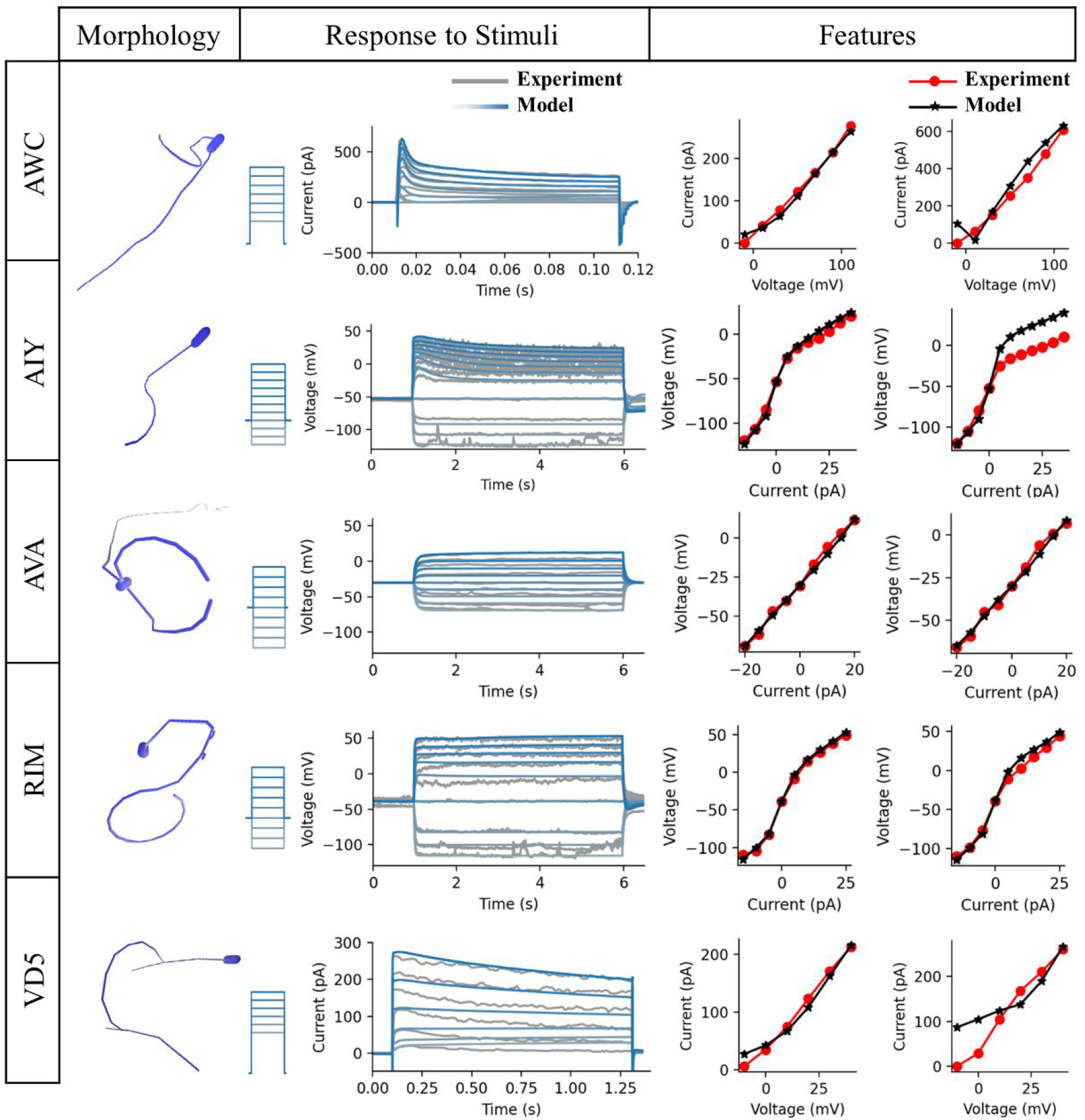
Models of five representative neurons fitted to electrophysiological recordings in experiments. AWC(L), AIY(L), AVA(L), RIM(L), VD5 were selected as the representative neurons for each of the five functional groups, respectively. Left, 3D morphology of each neuron^32,33^. Middle, membrane voltage/current dynamics induced by current or voltage clamp steps. The simulated responses (blue lines) are overlayed with biological experimental data (gray lines). The corresponding current or voltage clamp protocol are shown in left. AWC(L): voltage clamp protocol, in spans of 0.1 s, starting from -10 mV and increasing to 110 mV by 20 mV increments. AIY(L): current clamp protocol, in spans of 5 s, starting from -15 pA and increasing to 35 pA by 5 pA increments. AVA(L): current clamp protocol, in spans of 5 s, starting from -20 pA and increasing to 20 pA by 5 pA increments. RIM(L): current clamp protocol, in spans of 5 s, starting from -15 pA and increasing to 25 pA by 5 pA increments. VD5: voltage clamp protocol, in spans of 1.2 s, starting from -10 mV and increasing to 40 mV by 10 mV increments. Right, comparison between experimental (red dot) and simulated (black star) I-V curves, including I-V curve at steady-state (left) and I-V curve at the time when the rates of voltages/currents change reach a certain threshold (right). Experimental data are from published articles^36-39^.

These five neurons are representative for five different functional group: sensory neuron, interneuron, command neuron, head motor neuron and body motor neuron. They serve as the initial pool of digital neuron models to reconstruct other neurons in the neural network model. For neurons that are not representative neurons, their biophysical parameters were set the same as the representative neuron models that belonged to the same functional group. The referenced neurons of all neurons are listed in Supplementary Table S4. This approach allows us to achieve a high level of biophysical and morphological realism in our *C. elegans* neural network model with limited accessible experimental data.

For connections, we first modeled gap junctions as simple ohmic resistances, and synapses as continuously transmitting synapse, according to published models^32,40-43^. Then, we developed an algorithm to determine the number and locations of connections. The number of connections between two cells was from the cell adjacency matrix^10,11^ (Fig. 4A). For each connection, we randomly assigned a distance following the neurite centroids distance distributions of synapse/gap junction in experiment^13^ (Fig. 4B). Each connection linked two segments whose distance was closest to the randomly assigned distance. This approach allowed us to generate connections that were constrained by experimental connection numbers and distance distributions (Fig. 4C). Figure 4D-F show some examples of the connections between neurons in our neural network model. Most synapses and gap junctions were located on neurites that were relatively close to each other. Also, as shown in Fig. 4F, the constructed connections tended to cluster together, which followed the clustered organization principle in published research^13,14^.

**Fig 4.**
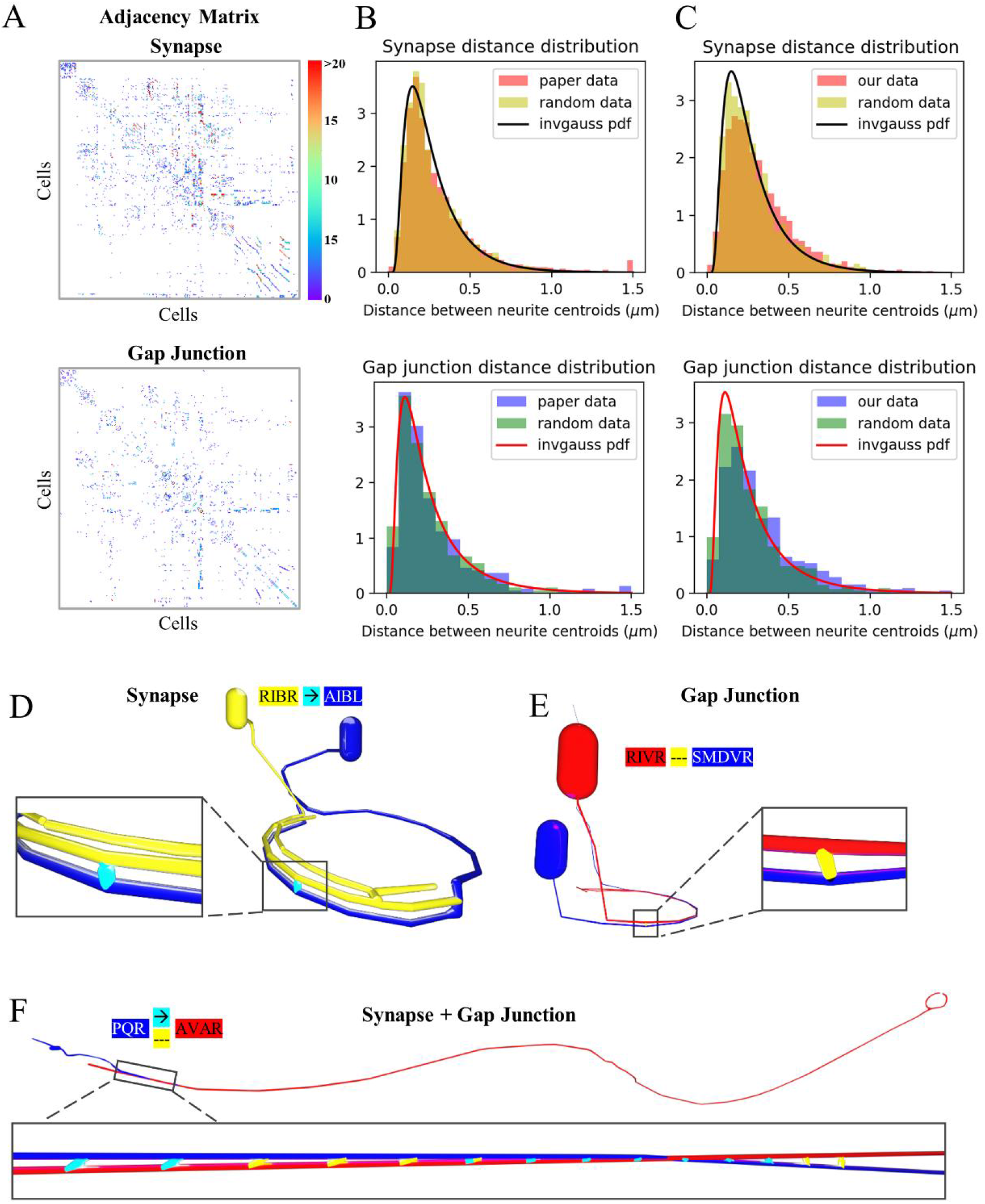
Connecting multi-compartmental neuron models in the neural network model. **A**. Synapse and gap junction adjacency matrices of the neurons from published data^10,11^. Matrix elements represent the total number of electron microscopy series capturing synapses (or gap junctions) between two neurons. The color indicates the series number. The order of cells in rows and columns corresponds to Supplementary Table S4. **B**. Distribution of distances between centroids coordinates of neurites at connected synapse (upper) and gap junction (lower) locations. Experimental data (red/blue bars) from a published paper^13^ is fitted by two inverse Gaussian distributions (black/red lines). Random data (yellow/green bars) is produced following the fitted inverse Gaussian distributions. **C**. Similar to B, with red and blue histograms indicating distributions in our neural network model constructed in our model. **D**. 3D visualization of a synapse (cyan) between RIBR (pre-synaptic, yellow) and AIBL (post-synaptic, blue). **E**. 3D visualization of a gap junction (yellow) between RIVR (red) and SMDVR (blue). **F**. 3D visualization of clusters of synapses (cyan) and gap junctions (yellow) between PQR (blue) and AVAR (red).

To determine the polarities of synapses (excitatory or inhibitory) and the connection weights, we employed a gradient-descend-based algorithm to minimize the mean squared error (MSE) between simulation results and the optimization target. In this study, our optimization target was a Pearson correlation matrix of 65 identified neurons in a prior study^24^, where activity series were recorded through whole-brain single-cell-resolution Ca^2+^ imaging. Figure 5A illustrates the correlation matrices generated from our simulation and the experimental data. These two maps exhibited similar correlation strengths, whose mean squared error was only 0.076.

**Fig 5.**
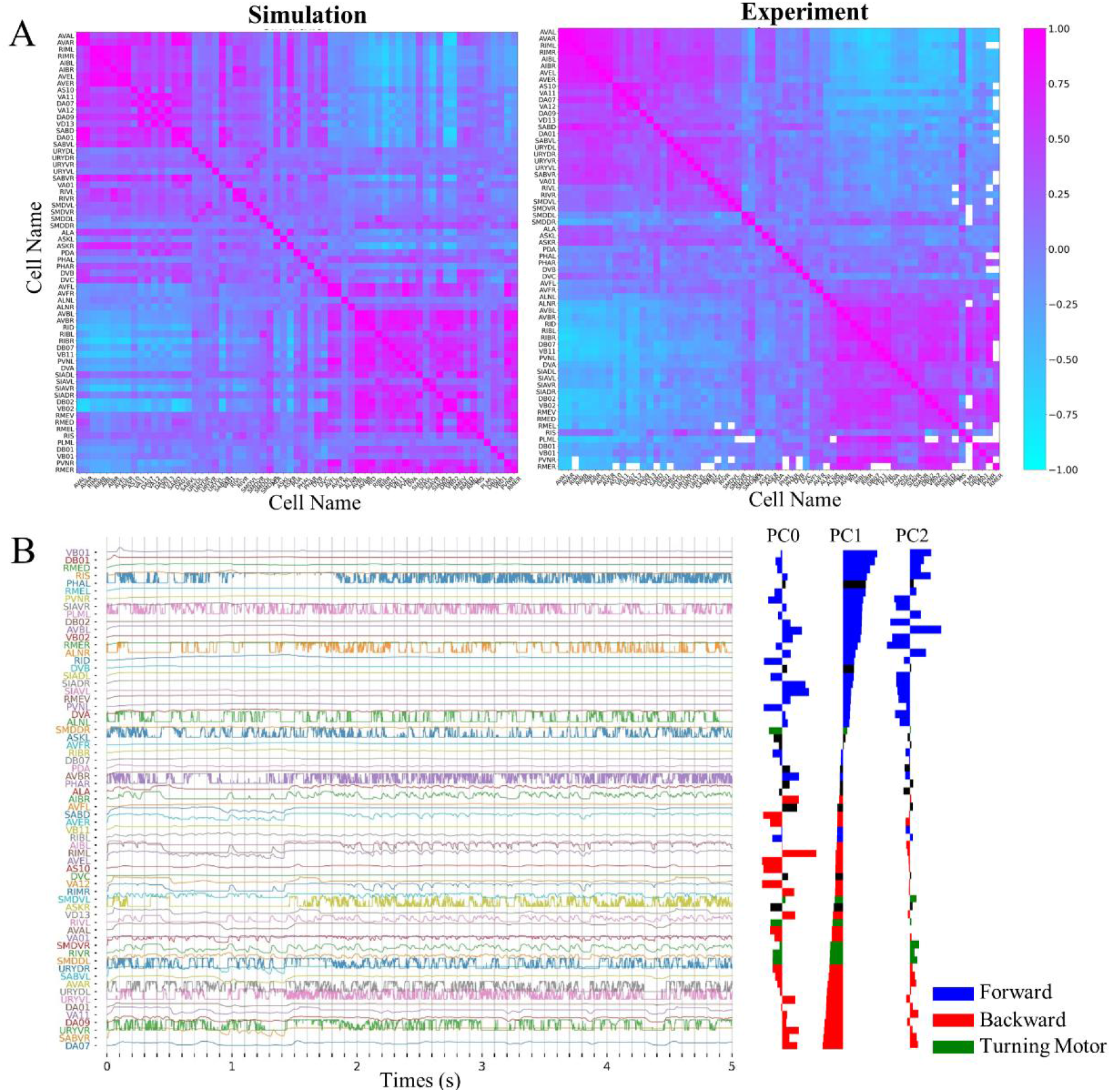
Neural network model of *C. elegans* fitted to functional map in biological experiment. **A**. Pearson correlation matrix of neurons in simulation and experiment^24^. The simulation matrix was derived from our optimized neural network model. Red and cyan color indicate positive and negative correlation coefficients, which were calculated by membrane potentials of two neurons. **B**. Membrane potentials of 65 neurons, each represented in a separate row. Principal component (PC0, PC1, PC2) weights of each neuron are shown by the bar plots on the right. Neurons were sorted by PC1 weights. Color of the bars indicates the reported role of the neuron in locomotion from published research^24^ (blue: forward neurons, red: backward neurons, green: turning motor neurons, black: no clear role).

To further analyze the optimized neural network model, we conducted principal components analysis (PCA) on the membrane potential of the 65 identified neurons, involving the calculation of principal components (PCs) based on the covariance structure in the normalized data (Fig. 5B). Our analysis unveiled that PC1 exhibits an opposite dynamic to that of PC2 (Fig. 5C). Upon sorting based on PC1, most neurons controlling forward locomotion (e.g., AVB, VB, DB neurons) and backward locomotion (e.g., AVA, VA, DA neurons) were distinctly classified into two groups (Fig. 5B). This indicates that our neural network model accurately captures realistic neural dynamical relationships, and the dynamics of neurons encode functions similar to those observed in real *C. elegans*, to some extent.

### Constructing a high-performance body & environment model of *C. elegans*

Next, we developed the body & environment model of the *C. elegans*. Our primary contributions are: 1) modelling the soft body, muscles, and solving soft body deformation, 2) creating the 3D simulation environment and solving soft body-fluid interaction, 3) proposed a novel reference coordinate system for behavior analysis. The framework of this model is illustrated in Figure 6A and B.

**Fig 6.**
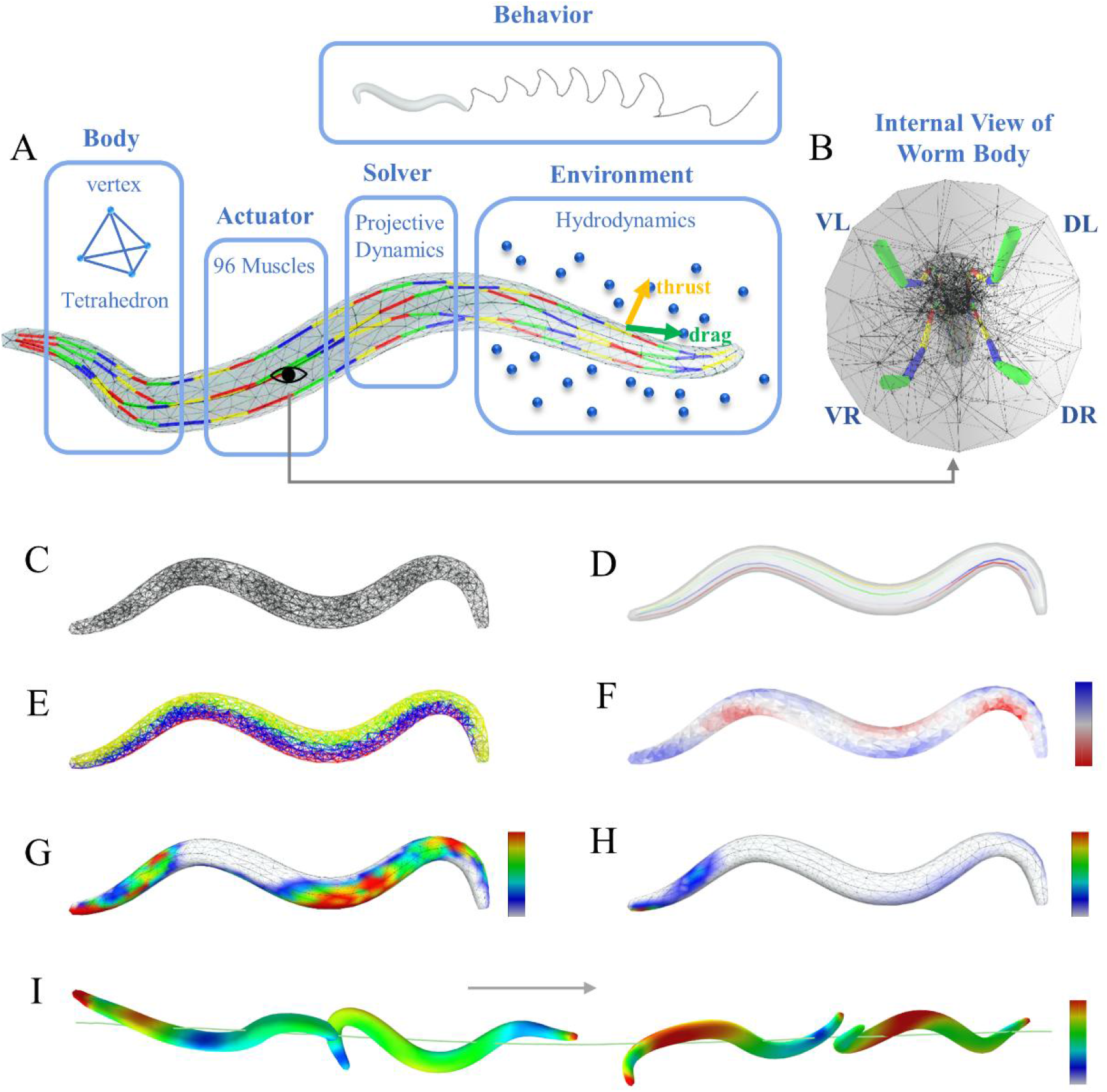
A biomechanical body model of *C. elegans*. **A**. Schematic diagram of *C. elegans* body model. The body was composed of tetrahedrons. 96 muscle cells along 4 muscle strings were serving as actuators for soft body deformation. Different colors on muscles indicate distinct muscle cells. The body was situated in a large 3D fluid environment and can move in the environment. Projective dynamics functions were used as the soft body solver, and simplified hydrodynamics calculated thrust force and drag force on the body surface. **B**. Internal view of the body model, consisting of 4 muscle strings: ventral right (VR), ventral left (VL), dorsal right (DR), dorsal left (DL). **C**. Triangulated tetrahedron mesh of the body. **D**. Four muscle strings: VR (green), VL (yellow), DR (red) and DL (blue). Color intensity reflects the strength of muscle contraction, with high intensity indicating contraction and transparent intensity indicating relaxation. **E**. Tetrahedrons driven by four muscle strings: VR (green), VL (yellow), DR (red) and DL (blue). **F**. Deformation of the soft body. Blue tetrahedrons are increase their volume, while red ones are decreasing. **G-H**. Thrust force (G) and drag force (H) on body surface. Red indicates strong force and blue indicates weak force. **I**. Positions and velocities of the body at four moments. The body was moving from left to right. Color indicates velocities, with red representing fast, blue representing slow.

The body model was built with tetrahedral mesh, including 984 vertices, 3341 tetrahedrons and 1466 triangles on the body surface (Fig. 6C). Also, it had four muscle strings: ventral right (VR), ventral left (VL), dorsal right (DR) and dorsal left (DL). Each string had 24 muscle cells arranged from head to tail (Fig. 6D). Tetrahedrons were controlled by their near muscle strings. The corresponding strings were visualized with different colors in Figure 6E. We used Finite Element Method (FEM) solver to compute positions and velocities of all vertices on the deformable body at any given time. In the computation, the solver used 4558 muscle constraints and 3341 corotated elasticity constraints. Thus, our soft-body simulation engaged a sum of 7,899 mechanical constraints. After the geometry construction, we need to solve the body deformation efficiently. In our model, we selected a real-time, highly robust, and accurate approach named projective dynamics as the FEM solver for calculation. The solved body state was visualized in Fig.6F.

We constructed a large 3D fluid simulation environment, where the body, foods were embedded. This environment was essentially a large cube, with each edge equating to approximately 1200 times the length of a *C. elegans*. The mechanic of the body movement in fluid is essentially a soft body-fluid coupling problem. To simplify this problem, we only calculated the thrust force (Fig. 6G) and drag force (Fig. 6H) on the body surface^44^. This simplification allowed for the simulation of a much larger space and improved computational efficiency. To achieve forward locomotion behavior, we used projective dynamics to calculate real-time bending waves that propagated from head to tail. The interaction between these bending waves and the fluid generated impetus for forward locomotion (Fig.6I).

*C. elegans* is an animal with soft body that exhibits special patterns during locomotion, wherein every point on its body oscillates continuously. Therefore, it is difficult to quantify the trajectory and body state during locomotion. Here, we proposed an effective and simple method to measure *C. elegans* locomotion. The main idea of this method is to establish a numerically stable coordinate system relative to the body itself, named Target Body Reference Coordinate System (TBRCS) (Fig. 7A). First, we defined the target body, standard body and Standard Body Coordinate System (SBCS). Then, we computed transformation M between the target body and the standard body. Finally, we used M to perform three-dimensional transformation on the SBCS to obtain the TBRCS. Details of this method are described in Methods. Using this method, we can stably quantify not only the body’s locomotion trajectory (Fig. 7A), but also the body states including the relative position and velocities of any points on the *C. elegans* body (Fig. 7B and Fig. 7C) and steering angle (Fig. 7D). For example, figure 7E displays the relative positions of 17 sampled 3D tracking points, while Figure 7F illustrates their corresponding velocities. In Figure 7E, the swing direction of body was Z direction of the TBRCS (dorsal-ventral direction). Also, there was a spring movement in X direction of the TBRCS (head-tail direction) during forward locomotion. Locomotion in Y direction of the TBRCS (left-right direction) was nearly flat since muscle activations were approximately symmetric in this direction. Figure 7F shows that, in this case, the primary propulsion of forward locomotion originated from tail (pink line), since speed represented power. Moreover, our method enables the quantification of the body’s steering. Figure 7D shows an example of a steering angle θ, which was defined as the angle between TBRCS x-axis and the direction of velocity. The steering angle θ was also numerically stable.

**Fig 7.**
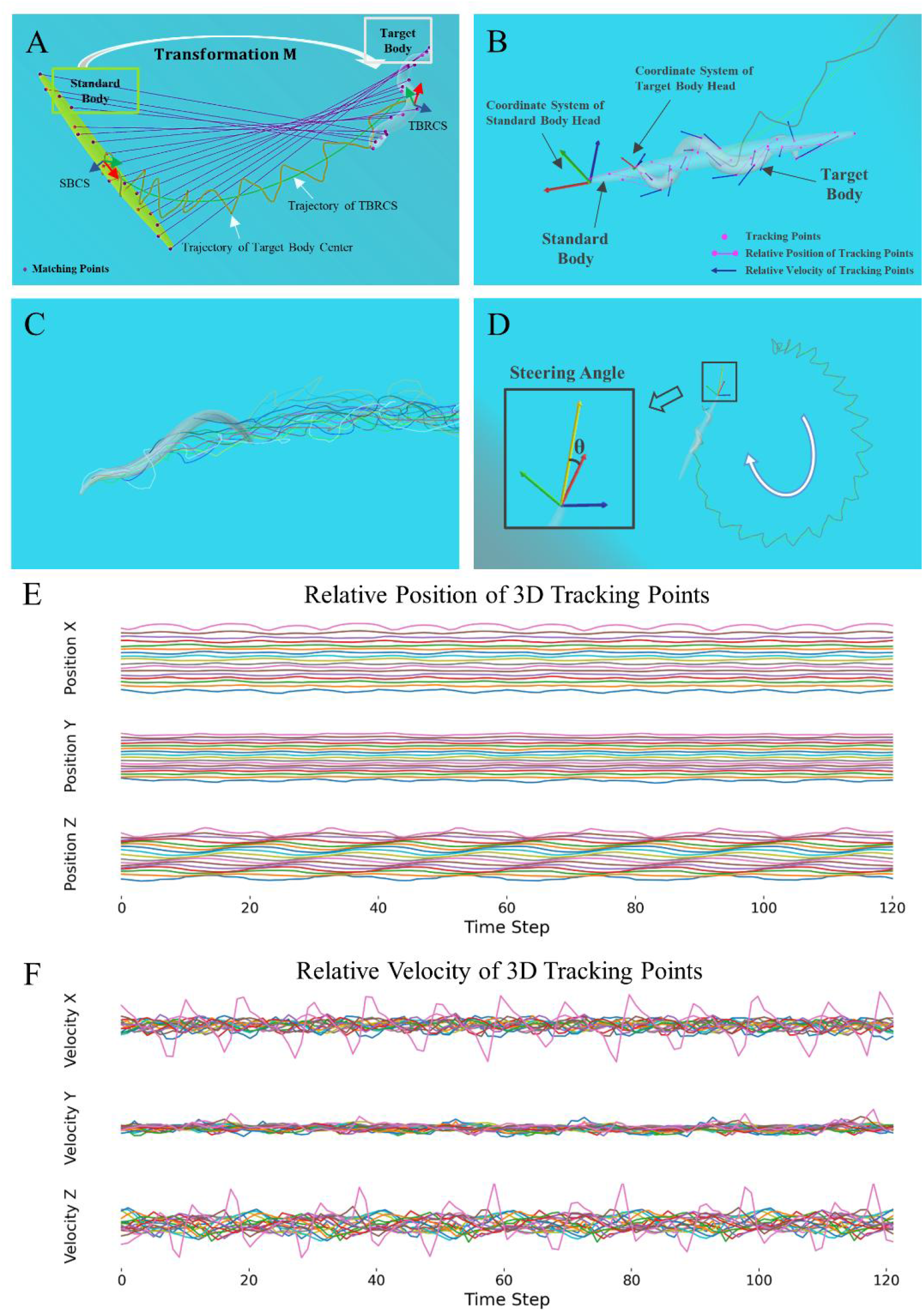
Quantification of body’s behavior. **A**. Schematic diagram of Target Body Reference Coordinate System (TBRCS) to quantify behavior stably. The standard body and SBCS are depicted on the left. The target body and TBRCS are illustrated on the right. Matching points (magenta dots) are positioned on both the standard and target body. Magenta lines links the corresponding points on two body. The transformation matrix M is obtained through singular value decomposition, enabling the conversion from SBCS to TBRCS. The trajectory of the target body center (orange line) exhibits a zigzag pattern, whereas the trajectory of TBRCS (green line) is smooth. **B**. Tracking body movement with TBRCS. 17 tracking points were defined on the surface of both the standard body and the target body. Magenta lines represent relative positions, while blue arrows indicate relative velocities. Coordinate systems of the head on standard body and target body are drawn out to display translation and rotation of the head on target body. **C**. Trajectories of 17 tracking points on the body. **D**. Steering angle θ of the body. The velocity vector is depicted as the yellow arrow, and the XYZ axes of TBRCS are represented by red-green-blue arrows. θ is defined as the angle between the red arrow and the yellow arrow and remains stable over time. **E-F**. Relative positions (E) and velocities (F) of 17 tracking points on the body during the movement.

In summary, our body model is simple yet preserving anatomical authenticity. Our environment model provides a large-scale 3D scene for simulations. Our simulation is real-time and allows simulations of *C. elegans* populations. Importantly, our model facilitates easy quantification of behaviors.

### Replicating locomotion behavior of *C. elegans*

In our model, we successfully replicated the locomotion behavior of *C. elegans* by a closed-loop interaction between the neural network model and the body & environment model. Specifically, the neural network model sensed the dynamic food concentration in the environment by sensory neurons. This sensory information was used to calculate the membrane potentials of motor neurons. The membrane potentials of motor neurons were subsequently translated into muscle activation signals, which controlled the movement of *C. elegans*’ body to navigate towards the food in the environment. An overview of this closed-loop simulation is illustrated in Figure 8A. Notably, we observed that the trajectory of the *C. elegans* movement in our model closely resembled the trajectory recorded in the experiments^45^, with both exhibiting a zigzag pattern (Fig. 8B). Additionally, *C. elegans* in both the simulations (Supplementary Movie S1 and Supplementary Movie S2) and experiments exhibited a dorsoventral fluctuation during 3D movement^46,47^.

**Fig 8.**
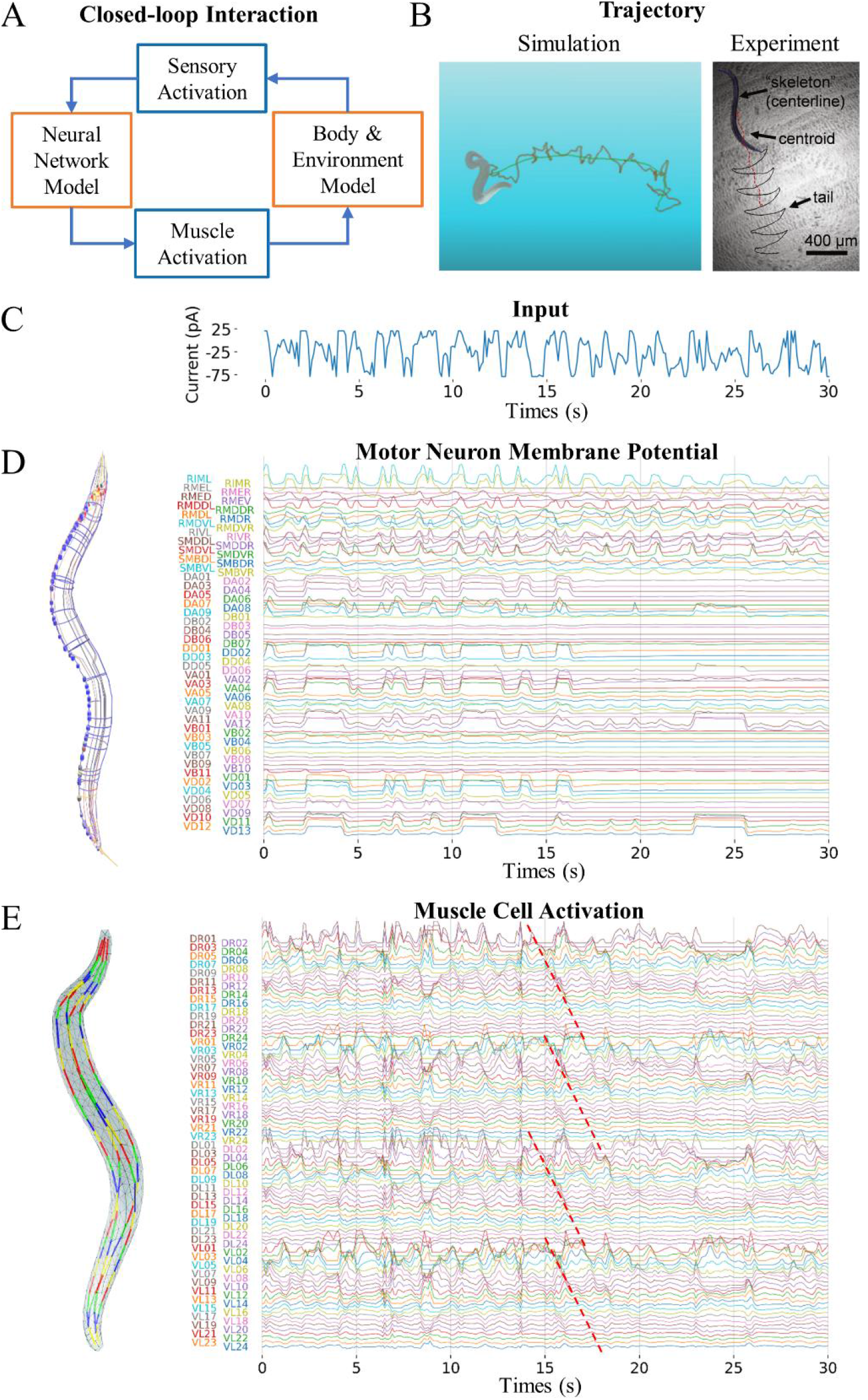
The closed-loop 3D simulation of *C. elegans* locomotion behavior. **A**. The information flow diagram of the closed-loop interaction between neural network model and body & environment model. **B**. Comparison of *C. elegans* trajectory during forward locomotion in simulation (left) and experiment (right, from a published article^45^). **C**. Input current injected to sensory neurons during locomotion. **D**. Left, neural network model of *C. elegans*. Right, membrane potentials of all motor neurons in the model during locomotion. **E**. Left, the body model with 96 muscle cells. Right, the activation signals of all muscle cells during locomotion. The dashed red line indicates that the activation signal shifting from head to tail.

Regarding sensation, we assumed that the food concentration in the environment follows a linearly distribution. To incorporate this sensory information into our model, we used the derivative of the food concentration as the amplitude of an external current injected into the soma of all sensory neurons. Regarding neuromuscular coupling, we employed a linear transformation, treating the neural network model as a reservoir to generate muscle activities^48^ (Methods 3.6).

During locomotion, the input of sensory neurons exhibited fluctuations due to the zigzag movement of the body’s head (Fig. 8C). The membrane potentials of each individual motor neuron also oscillated in response to the sensory input, especially the head motor neurons (Fig. 8D). The activation of muscle cells revealed traveling waves from the head to the tail, accompanied by alternating contractions and relaxations of dorsal and ventral muscle cells (Fig. 8E). These findings closely resemble observations in biological experiments^49^. To facilitate visualization, we have incorporated the locomotion of *C. elegans*, the activities of its neural network model, and muscle activities into our graphical user interface (GUI) (Fig. 9).

**Fig 9.**
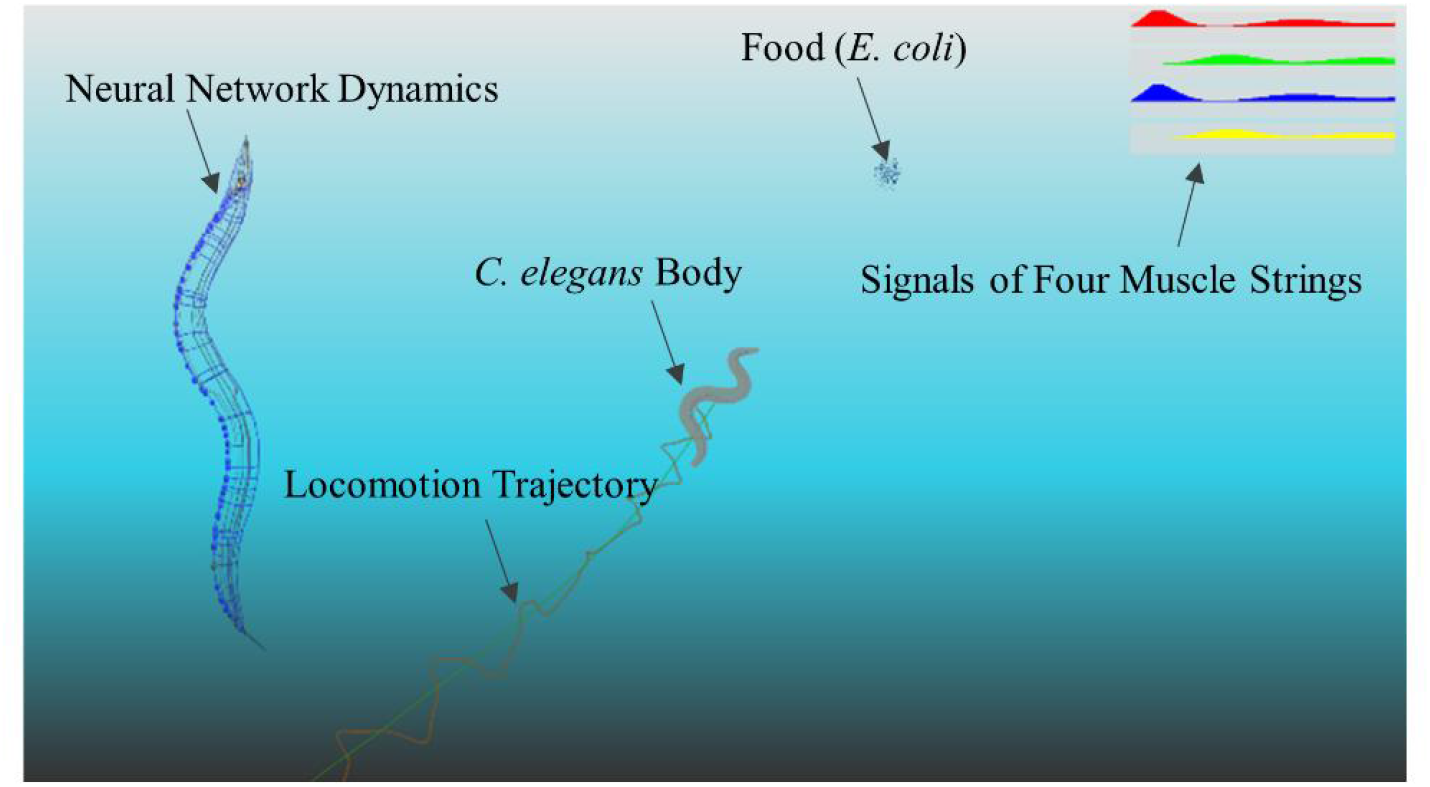
Graphical User Interface (GUI) of MetaWorm. Graphical User Interface (GUI) of MetaWorm, including: the neural network of *C. elegans* where the color indicates membrane potentials, a *C. elegans* body actively moving towards a food source (*E. coli* population) in the 3D fluid environment, the locomotion trajectory, activations of four muscle strings extending from head to tail.

In summary, our model successfully reproduced locomotion behavior in this closed-loop simulation and facilitated the visualization and analysis of neuron dynamics, muscle activities, and behavioral patterns.

### Exploring the impact of neural network structure on dynamics and behaviors

We are interested in exploring the extent to which network structure influences the neural dynamics and behaviors, thus we conducted synthetic perturbations on the structure of our neural network model. Synthetic perturbations of the neural system often exceed individual tolerance level in animals. However, our model enables researchers to carry out experiments without concerns about experimental technique limitations or animal tolerance. Based on MetaWorm, we performed experiments involving synthetic perturbations to quantitatively examine the impact of network structure on network dynamics and behaviors.

In the absence of neurites, the body exhibits a higher degree of twisting during locomotion, as indicated by the larger relative locations of tracking points on the head and tail compared to the control group. This suggests that neurites play a crucial role in the computations associated with network information processing (Fig. 10B). Moreover, shuffling connection locations within the model leads to faster oscillations of the head and tail but slower forward locomotion (Fig. 10C). This suggests that the specific locations of connections along the neurites are important, potentially facilitating localized computations within neurons. Additionally, despite the neural network model contains fewer gap junctions than synapses, the role of gap junctions in *C. elegans* neural network appears to be more important than that of synapses (Fig. 10D-G), because perturbations on gap junctions result in more obvious changes in the Pearson Correlation matrix compared to perturbations on synapses. Perturbations on gap junctions accelerate the twisting of the body compared to the control group, whereas perturbations on synapses result in slower twisting. These results emphasize the importance of network structure in shaping network dynamics and behavior patterns. By utilizing our model, researchers can gain valuable insights into the contributions of network structure to neural network.

**Fig 10.**
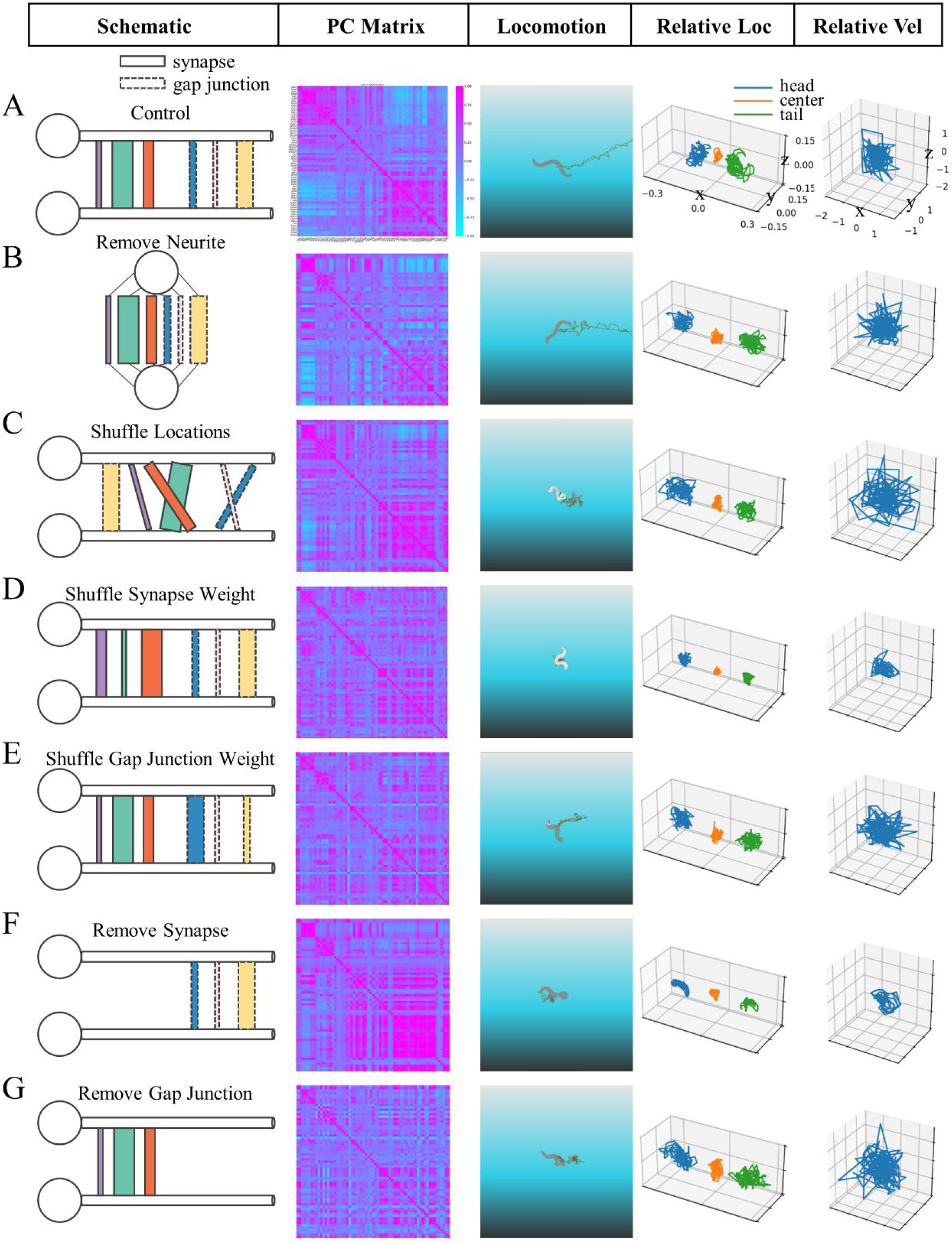
Synthetic perturbation of on the neural network model in MetaWorm. **A**. Column 1: Schematic diagram illustrates connections between two neurons in the neural network model. Ball and stick denote soma and neurite, respectively. Rectangles with solid and dashed lines indicate synapses and gap junctions. Rectangles with different colors represent different connections. The width of rectangles signifies connection weights. The depicted connections between this neuron pair represent all connections in the neural network model. Column 2: Pearson correlation matrix of neurons during simulation of the neural network model. Column 3: Locomotion behavior of the *C. elegans* body controlled by the neural network model in Column 1. Column 4: Relative position of 3D tracking points on head (blue), center (orange) and tail (green) of the *C. elegans* body. Column 5: Relative velocity of a 3D tracking point on head of *C. elegans* body. **B-G**. Diagrams represented as in **A**, but with different perturbation types: (**B**) Removal of neurites from all neurons in the neural network model, relocating connections onto the soma, (**C)** shuffling the location of connections on neurites while fixing source and target neurons, (**D)** shuffling the weights of synapses while fixing their locations, (**E)** shuffling the weights of gap junctions while fixing their locations, (**F)** removal of all synapses in the neural network model, (**G)** removal of all gap junctions in the neural network model. These perturbations are sequenced based on their distance to the control group in terms of the Pearson Correlation matrix (B: closest, G: farthest).

## Discussion

In this study, we present MetaWorm, a data-driven model of *C. elegans* that integrates models of neural network, body and environment. The neural network model of MetaWorm is highly detailed and data-driven, capturing neural population activity patterns and encoding functions observed in real *C. elegans*. The body & environment model of MetaWorm is realistic, efficient, and allows for the quantification of behaviors. Based on the interaction of these two sub-models, we achieved locomotion behavior in *C. elegans*. This represents the first time that an integrative data-driven model replays a behavior with motor control and sensory feedback.

The neural network model of MetaWorm integrated experimental data including ion channel dynamics, neural morphologies, electrophysiology, connectome, synaptic organization rules on neurites, as well as neural activities. Although we collected and integrated as much data as possible up until 2023, there remained a substantial amount of detail that could not be fully implemented due to lack of data. Also, the physical parameters in body & environment model did not correspond precisely to those of actual *C. elegans*, because they were difficult to measure accurately. However, with the influx of new experimental data, these models can be enhanced and refined.

For the neuromechanical connections from motor neuron to muscle, the neural network model served as a reservoir that processed data in a complex and nonlinear way. The input was mapped onto an output linear layer to produce muscle activation. With only one linear readout layer, the neural network model successfully controlled the body’s locomotion. This demonstrates the great information processing ability of our neural network model. In the future, more realistic models of the neuromuscular junction can be used in our neural network model, reflecting the real synaptic properties from motor neuron to muscle.

When the input of the neural network model was a periodic standard signal, i.e. without sensory feedback from the environment, the membrane potentials of motor neurons and the activations of muscle cells were periodic (Supplementary Figure S1). This resulted in the movement trajectory of the *C. elegans* body following a standard Z-shape, which was extremely similar to the trajectory in the experiment^45,50^ (Supplementary Figure S1B and Supplementary Movie S3). Conversely, When the input was a sensory signal dependent on the body’s location in the environment (Fig. 8), the input was not periodic, leading to non-periodic dynamics in both neurons and muscle cells. Therefore, the trajectory of the body was not as regular as that observed in the experiment (Fig. 8B). Future studies will focus on designing more reasonable Central Pattern Generators (CPGs) or motor control systems to achieve more realistic movement.

Training biophysically detailed neural network model is challenging and consumes much time and GPU memory resources. In the future, we will improve the optimization algorithm to effectively train larger detailed neural network that contains all 302 neurons, with more constrains such as single neuron voltages, and reproduce more behaviors beyond zigzag locomotion.

MetaWorm serves as a white box model of *C. elegans*. Its two sub-models and their interaction can be independently modified or extended. We provide researchers with user-friendly interfaces to design their own model based on their experimental data. Moreover, our model serves as a valuable reference for designing integrative data-driven models for other animals. In this era of exponential growth in experimental data, these integrative data-driven models will enable us to gain a deeper understanding of the neural computation that underlies specific behaviors.

## Acknowledgements

Funding: this research was funded by National Key R&D Program of China (2020AAA0105200)

## Author Information

### Contributions

M.D.Z. developed the neural network model module, designed and conducted experiments, and interpreted results. N.W. developed the body & environment module. H.X.M. and L.M. developed the 3D neural visualization software. X.R.J. and M.D.Z. collected experimental data for constructing the neural network model. X.Y.M. and M.D.Z. optimized the connection algorithm and rewrote the code of the neural network model module for open source. G.H. developed the optimization algorithm for the multi-compartmental neural network model. M.D.Z., N.W., and X.R.J. wrote the article. K.D. reviewed the research and provided advice. L.M. supervised the research, coordinated and conceptualized the study, and contributed to writing the article. All authors participated in the paper revision process.

### Corresponding authors

Correspondence to Lei Ma

## Ethics declarations

The authors declare no competing interests.

## Methods

### 1 Details of Neural Network Model

#### 1.1 Neuron models

Neurons were modeled by morphologically derived multi-compartmental models with somatic Hodgkin-Huxley dynamics^1^ and passive neurites. These neuron models captured both the biophysical properties and spatial complexity of neurons. The models integrated three kinds of neuron data: ion channel, morphology, and electrophysiology.

Ion channel models were determined according to published studies^2^. There were 14 ion channels divided into 4 categories: voltage-gated potassium channels (SHK-1, EGL-36, SHL-1, KVS-1, KQT-3, EGL-2, IRK1/3), calcium-regulated potassium channels (SLO1, SLO2, KCNL), voltage-gated calcium channels (EGL19, UNC2, CCA1), and passive sodium channel (NCA). Model equations and parameters for ion channels are given in Supplementary Table S1 and Supplementary Table S2.

As for morphologies of neuron models, the spatial structures were adopted from the *C. elegans* neuronal anatomy models created by the Virtual Worm Project^3^ and the OpenWorm Project^4^. Each neuron model had a soma section and several neurite sections. Neurite sections were further divided into several compartments, each of which was shorter than 2 μm.

Only soma compartment had active ion currents. Soma compartments and neurite compartments shared the same passive properties. To identify the passive and active parameters of neuron models, a deep learning tool^5^, was used to fine-tune these parameters to reproduce experimental data of voltage/current clamp. The value of the parameters can be found in Supplement Table S3.

Less than 20% of the *C. elegans* neurons have been recorded for the desired electrophysiology data. To estimate model parameters for those neurons that were not recorded, a method was implemented to generalize parameter values. First, considering the available data and hierarchy of information flow in *C. elegans* nervous system^6,7^, the neurons were divided into 5 functional groups (sensory neuron, interneuron, command interneuron, head motor neuron, and body motor neuron, see Supplement Table S4). Second, 5 neurons of each functional group (AWA^8^, AIY^8^, AVA^9^, RIM^8^, and VD5^10^) were selected and their model parameters were optimized as described above. These parameter values of optimized neuron models were then generalized to other neuron models according to their functional categories. In other words, neuron models in the same functional group share identical electrical parameters. Each neuron models have their own morphologies yet similar electrical properties to those neurons in the same functional group.

#### 1.2 Graded synapse and gap junction models

The neuron models were connected by graded synapses^8,11^ and gap junctions^12^ in *C. elegans* neural network. Graded synapses, include excitatory and inhibitory synapses, were modeled according to published *C. elegans* models^4,13,14^. The synaptic conductance is continuously changed with presynaptic membrane potential. Postsynaptic current is given by:

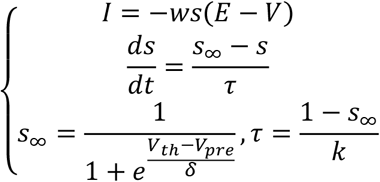

where *w* is maximal postsynaptic membrane conductance for the synapse, *s* is a voltage-sensitive parameter controlling the conductance, *E* is the reversal potential, *τ* is the time constant of s, *V*_*th*_ is the presynaptic equilibrium potential at the middle of the voltage sensitive range, *V*_*pre*_ is the pre-synaptic membrane potential.

Gap junctions were modeled as simple resistances, where current flowing from presynaptic cell to postsynaptic cell is given by:

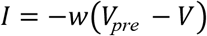

where *w* is the conductance of gap junction.

The parameter values of these synapses were set according to published models^12^: the conductance of the synapses and gap junctions had the same order of magnitude of 1e-4 μS. The specific weight of each synaptic connection in a specific neural network can be set randomly or be optimized according to a target, as described in Method 1.4.

#### 1.3 Connection locations

Detailed neuron connections in the neural network model were located on the neuronal neurite (axons or dendrites) of the neurons, which were based on the neuron connectivity matrices and a neurite connectivity algorithm.

The neuron connectivity matrices were acquired from published connectome data^6,15^. The original data in the matrices was the total number of electron microscopy (EM) series of all synapses (or gap junctions) between two neurons. In our work, the number of EM series was transformed into the number of synapses (or gap junctions) by:

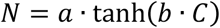

where *C* is the number of EM series, *a* and *b* are constants. For synapses number transformation, *a =* 23.91,*b =* 0.02285, while for gap junction, *a =* 20.49, *b =* 0.02184. These parameters were inferred from partial-animal connectome data^6^.

After the number of synapses (or gap junctions) was determined, the locations of synapses (or gap junction) on neural neurites were identified by a connection algorithm. The algorithm was based on the rule that distances between the centroid coordinates of possible presynaptic and postsynaptic neurites follow a certain statistical distribution^16^. We fitted synapse and gap junction distribution by two Inverse Gaussian distributions:

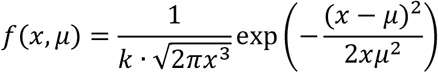

where *µ* = 0.44, *k* = 0.63 for synapse distribution, *µ* = 0.70, *k* = 0.40 for gap junction distribution. The algorithm of generating connection locations was designed:

##### Algorithm 1.

Generate connection locations.

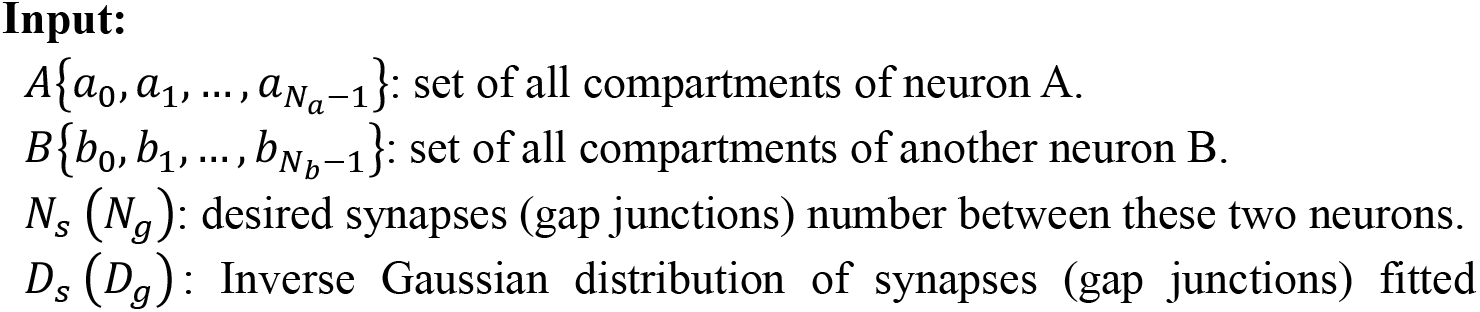
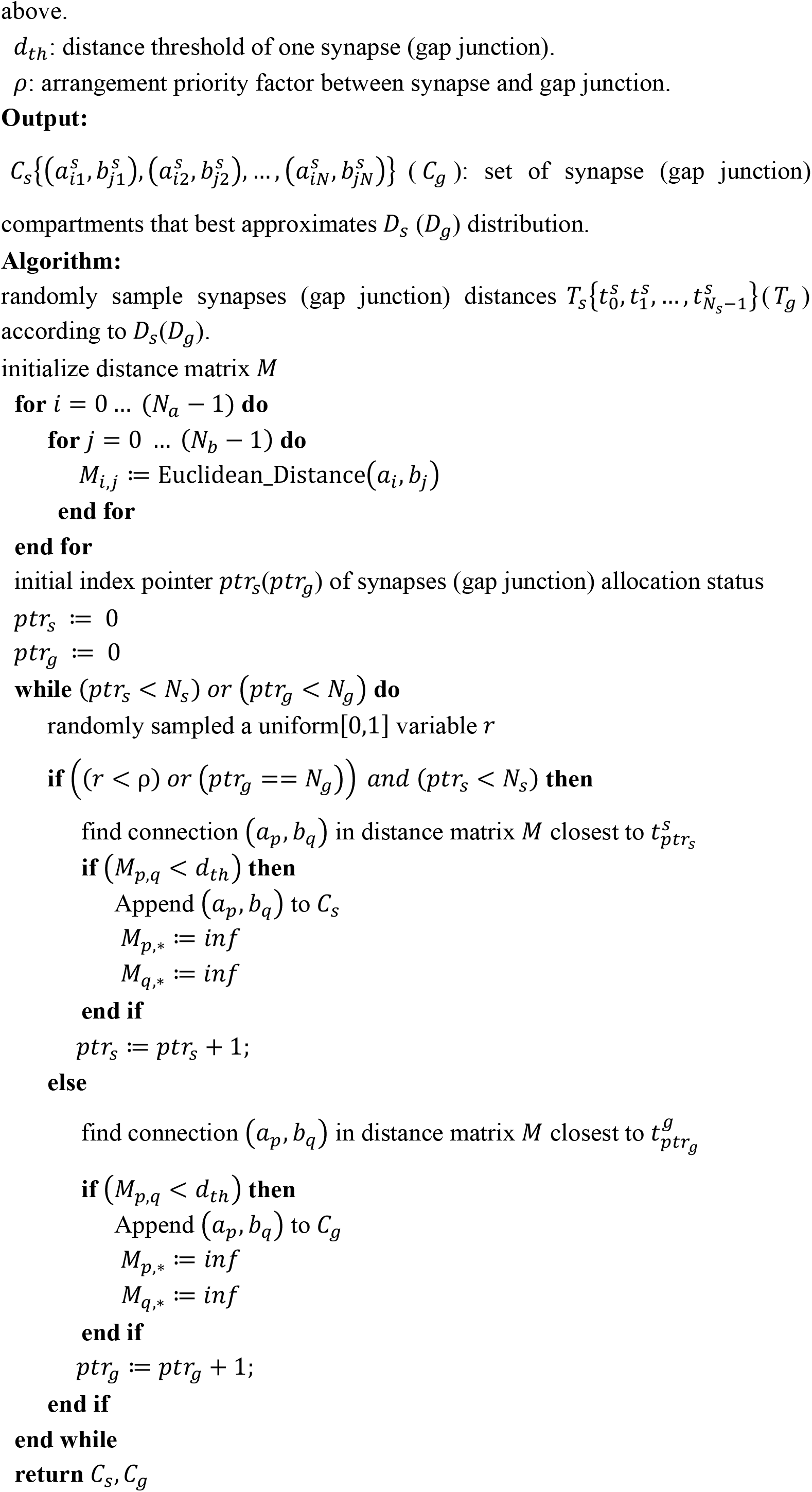

#### 1.4 *C. elegans* neural network model

Our neural network model consists of 136 neurons, including 15 sensory neurons for inputs and 80 motor neurons for outputs (Supplement Table S5). Here are the reasons why we chose these neurons. First, 65 head ganglia neurons, whose neural dynamics were recorded in the published whole-brain Ca^2+^ imaging experiment^17^ should be selected to enable the comparison between experiment and simulation. Second, other 71 neurons, most of which were motor neurons in the locomotion circuit of *C. elegans*^7,18^, were added to the network so that the model could be complete in a sensory-motor functional structure. In summary, this model contained neurons for chemosensory, decision, and locomotion.

#### 1.5 Connection parameters optimization

The weights of synapses and gap junctions in the neural network was initialized randomly. The polarities of synapses were initialized randomly according to a given excitatory/inhibitory ratio (2.33). These parameters were optimized using an unpublished optimization algorithm for multi-compartment neural network. The algorithm employed a mathematical approximation approach to calculate gradients for Back Propagation Through Time (BPTT) to update parameters. Except for the weights (of synapses and gap junctions) and the polarities (of synapses), the inputs (of some sensory neurons) also can be optimized by this optimization algorithm.

#### 1.6 PCA of neural activities

Principal Component Analysis (PCA) was applied to the time series data of soma membrane potential of the 65 head neurons. Before conducting the analysis, the time series data was preprocessed by subtracting the mean value of the trace.

#### 1.7 Correlation analysis

The correlation between two neurons was quantified by the Pearson correlation coefficient of their neural activities, which ranged from -1 to 1. A coefficient of -1 or 1 indicates a strong negative or positive correlation of neural activity, respectively. A coefficient of 0 indicates that the neural activities of the two neurons are unrelated. In this analysis, the neural activity was represented by the membrane potential of soma.

### 2 Details of Body & Environment Model

#### 2.1 The body model

Tetrahedron is the simplest ordinary convex polyhedra in three-dimensional space. It has four vertices, six edges, and is bounded by four triangular faces. We used tetrahedron as the basic element to construct the body model. Surface shape of the body was extracted as triangle list from Sibernetic’ membrane data^19^. Then we triangulated the body with a robust and high-quality algorithm^20^. The triangulated tetrahedron mesh was refined for latter mechanics solver. The final body mesh was in a straight, natural resting pose that we defined as the standard body (Fig.6A). The triangulated tetrahedron body was not strictly symmetric in dorsal-ventral direction and left-right direction. Since triangulation algorithm had additional constraints on quality of tetrahedrons. This non-strict symmetry is probably more consistent with real *C. elegans*.

#### 2.2 Muscles

96 muscles were arranged from head to tail according to *C. elegans* anatomy. Each 24 muscles were grouped into longitudinal string, marked as ventral right (VR), ventral left (VL), dorsal right (DR), dorsal left (DL) in four quadrants (Fig.6B)^19^. We imported body membrane mesh into blender^21^ and modeled VR, VL, DR, DL by four Bezier curves. Then muscle geometry was exported as wavefront obj file. We used Finite Element Method (FEM) to simulate the soft body, and used finite element constraints to simulate elastic muscle cells of body. Each muscle cell (Fig.6A) corresponds to one or more FEM constraints according to the discretized tetrahedron body (Fig.6C). Only nearby tetrahedrons can be driven by corresponding muscle cells.

#### 2.3 Soft body solver

We used projective dynamics^22^ as the solver of the soft body. The purpose of mechanics solver is calculating positions and velocities of all 3D vertices of the deformed body over time. Key idea of projective dynamics is to compute a local step and a global step iteratively. At local step, vertex positions of the body are fixed and projection variables are updated. At global step, projections are fixed and vertex positions of the body are updated. The local step is well suited for parallelism, thus can be optimized for performance. The global step is a linear system and can be precomputed if the body geometry and constraints are not changed. After a few local-global iterations the vertex positions of deformed body are converged. Then velocity of deformed body can be computed by Euler integration.

#### 2.4 Interaction with environment

A simplified model of hydrodynamics was used to simulate the interaction between the body and the fluid, similar to the published study^23^. For simplicity, we supposed external force is only formed at the surface of body. Two kinds of external forces were considered here: thrust force offered by body movement, and drag force offered by fluid. With this simplification, computation in the whole 3D fluid environment is avoided. Thus, more agents, larger environments, more complex tasks are achievable through this simplification.

#### 2.5 Numerical stable coordinate system for body movement

*C. elegans* is a soft body animal which has special patterns during movement. Every position of the body is oscillating during movement. Moreover, there is no apparent important marks on the body during locomotion, therefore it’s hard to find an apparent stable coordinate system for *C. elegans* movement. Simply using head or body center as a coordinate system can’t analysis the body’s movement quantitatively and stably. Here we proposed an effective and simple method to measure *C. elegans* locomotion. The core of our method is to find a numerically stable coordinate system relative to the body itself. The main steps of our method are as follows:

1. Define the target body, standard body and standard body coordinate system (SBCS).
2. Obtain transformation between the target body and the standard body.
3. Use the transformation in 2) to perform three-dimensional transformation on the SBCS to obtain the target body reference coordinate system (TBRCS).

Details of the three steps are described below:

Step 1) The target body is the research subjects. It can change its shape over time and exhibits various behaviors. The standard body is a special case of the target body. Here we define the standard body in a natural state where the head and tail are in a straight line, without stretching and bending. There is no internal force in the standard body soft body, and no external force acting on the standard body surface. The SBCS is a three-dimensional orthogonal coordinate system used to reflect the overall structure and function of the standard body. In this case, we define the SBCS with origin at the standard body center. X-axis of the SBCS is towards the head of the standard body. Z-axis of the SBCS is towards ventral to dorsal direction of the standard body. Three axes of the SBCS form a right-handed coordinate system. Schematic diagram of the target body, the standard body and the SBCS can been seen in Figure. 7A.

Step 2) In this step, we first define the corresponding matching points on the target body and the standard body. For example, simply use all the vertices of the body. The transformation between the matching points on the target body and the standard body is calculated as the transformation between the target body and the standard body. Calculation formulas of the transformation are as follows:

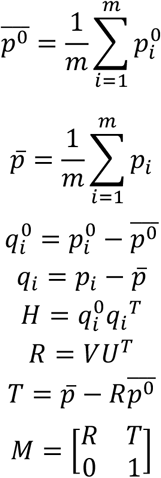

The matching points are represented by column vectors. 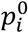 are matching points on the standard body, and *p*_*i*_ are matching points on the target body. m is the number of matching points. 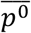 is the center point of matching points on the standard body, and 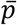 is the center point of matching points on the standard body. 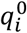 refers to the standard body matching points with the center offset subtracted, and *q*_*i*_ denotes the standard body matching points with the center offset subtracted. *H* is a matrix computed by 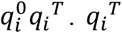 is transpose of 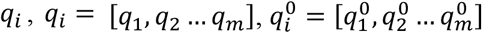.*U* and *V* are obtained by the singular value decomposition *H = U*Λ*V* of the matrix *H. U*^*T*^ is transpose of *U. M* is the transformation matrix between the target body and the standard body, which will be used in next step. *R* is rotation part of *M, T* is translation part of *M*. A schematic diagram of matching points and transformation *M* are shown in Fig. 7A.

Step 3) In this step, we use matrix *M* to perform three-dimensional transformation on the SBCS to obtain the TBRCS. The matrix of the TBRCS can be formulated as: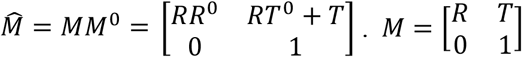, *R* and *T* have been described in last section.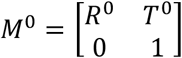 is matrix used to represent the SBCS. *T*^0^ is the origin of the SBCS, and *R*^0^ indicates orientation of the SBCS. The column vector of *R*^0^ represents the orthogonal coordinate axis. Finally, we can make the TBRCS through 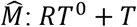 as the origin and *RR*^0^ as the orthogonal coordinate axis.

#### 2.6 Numerically stable measurements of body movement

##### 1) Measurement of body trajectory

During behavior analysis, *C. elegans* can be seen as a deformable biology composed of multiple points. When studying the locomotion of a deformable body, generally one or more points of biology body is defined as tracking object, and locomotion of deformable body is studied through tracking these defined points. For deformable biology like *C. elegans*, position and velocity of local tracking points are hard to reflect overall state of body locomotion (Fig. 7C). As a result, values such as body trajectory or steering angle obtained through these tracking points are usually unstable over time. Using the TBRCS, we can better describe the overall trend of body movement. Comparison of trajectories are shown in Figure. 7A. If the center point of body is used as trajectory, the trajectory is a zigzag shape. However, if the TBRCS is used as trajectory of body movement, the trajectory is a smooth curve which means stable values. It can be seen the TBRCS is more suitable as a measurement of overall path and direction of body locomotion.

##### 2) Tracking body movement

Here, we demonstrate how to track body movement with the TBRCS. If the standard body is drawn in the TBRCS, then it can be used as a reference for the target body. The standard body and the target body will be center alignment, translate and rotate together. We define 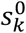 as 3D tracking points of the standard body, and s_*k*_ as 3D tracking points of the target body. k *=* 1, 2 … n. n is the number of tracking point. Then relative movement of body can be defined as 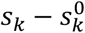. It’s a measurement of body locomotion relative to itself (the standard body), which is more numerically stable than relative to world. 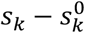 is also better than relative to body center, since both translation part and rotation part of whole deformation body are considered in the TBRCS. Moreover, sampling bias of tracking points is eliminated through subtract operation, since bias minus bias equals zero.

In Figure 7B, we sampled 17 tracking points on the body uniformly. Offset of the tracking points are indicated by pink lines between tracking points. Relative velocity of 3D tracking points is drawn in blue arrows. As a common method, trajectories of sampled 17 tracking points that relate to world are also shown in Figure 7C. We can see that compared to all 17 tracking points trajectories, trajectory of the TBRCS is the smoothest. Relative position and relative velocity of 3D tracking points over time are plotted in Figure 7E and Figure 7F respectively. In Figure 7E, Positions of 3D tracking points are tiled from bottom to top for clear seeing. In this case, swing direction (Z direction of the TBRCS) of swimming body can be clearly reflected with the TBRCS. We can also find that there is a spring movement in X direction of the TBRCS (tail to head) during forward swimming. Locomotion in Y direction of the TBRCS is nearly flat since in this case muscle activations are approximately symmetrical in this direction. From Figure 7F we can further deduced that the biggest propulsion of forward swimming is come from tail (purple line), since speed means power.

##### 3) Steering angle of body movement

If 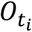 is the trajectory of TBRCS origin at time *t*_*i*_ · *t*_*i*_ *= t*_1_, *t*_2_ … *t*_*m*._Then we can get velocity of TBRCS by 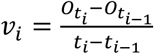. In this way, velocity magnitude and velocity direction over time are numerically stable. We used angle θ which is between TBRCS x-axis and *v*_*i*_ as a measurement of steering, as shown in Figure 7D. Both *v*_*i*_ and the TBRCS are numerically stable over time, thus θ is also numerically stable.

### 3 Locomotion Behavior Simulation of *C. elegans*

#### 3.1 Simulation configuration

The neural network simulation was coupled with the physical body & environment simulation to achieve closed-loop interaction. Food was at the center of the simulated physical environment, acting as an attractor to the *C. elegans*. The food concentration was linearly distributed in the environment. The neural system received currents regarded as sensory signals. The amplitude of the currents was proportional to the gradient of food concentration. The outputs of the neural network controlled the movements of the body in the environment through a reservoir readout weight matrix (see Method 3.2). As the body moved in the physical environment, the distance between the body and the food changed, causing the food concentration gradient updated and transformed to currents injected into the soma of sensory neurons.

During the interaction of the two sub-models, the simulation time step of the neural network was 5/3 ms and that of physical environment simulation was 100 ms. The two systems synchronized every 100 ms to update the sensory signals and muscle forces.

#### 3.2 Reservoir configuration

To establish the neuron-muscle connection between the neural system and physical body muscles, a reservoir computing framework was used. The neural network model acted as the non-linear reservoir. The membrane potentials of 80 motor neurons (Supplement Table S5) were linearly combined and mapped to 96 muscle forces via an 80 by 96 matrix.

The reservoir was trained using data extracted from a 10-second movement generated in the simulated physical environment. The pairs of food concentration and muscle force during the movements were used as inputs to the neural network and target outputs of the readout layer, respectively. Only the reservoir readout matrix was optimized, while the parameters in the neural network model was fixed.

### 4 User-friendly Interfaces

The neural network model consists of ion channel, neuron, and network connection modules. The parameter and algorithm settings of each module can be modified independently. Parameters were saved in JSON files that are easy to read and modify.

## Code Availability

The *C. elegans* neural network simulation was performed with the code written in Python 3.7 and employing NEURON 8.0.0^24^ as a simulation engine. The custom-developed code used in this work will be made publicly available in an online repository latest upon journal publication. If you wish to obtain the code before then, please contact us. To facilitate better evaluation of this article by reviewers, we also provide the code that can be run (we will continue to optimize our code details): https://www.dropbox.com/s/ep2yv3rmlp67eex/worm-simulation-share.zip?dl=0, Please contact us if you would like to have access to the latest code before our article is published.

