## Supplementary for "MetaWorm: An Integrative Data-Driven Model Simulating *C. elegans* Brain, Body and Environment Interactions"

### Supplementary Materials

##### Table S1. Ion Channel Models

| Ion Channel | Dynamic Model |
|---|---|
| **SHL1** | $m_{\mathrm{SHL1},\infty}(V) = \frac{1}{1 + e^{\frac{-(V-V_{0.5})}{k_a}}}$<br>$\tau_{m_{\mathrm{SHL1}}}(V) = \frac{a}{e^{\frac{-(V-b)}{c}} + e^{\frac{(V-d)}{\tilde{e}}}} + f$<br>$h^{\mathrm{f}}_{\mathrm{SHL1},\infty}(V) = h^{\mathrm{s}}_{\mathrm{SHL1},\infty}(V) = \frac{1}{1 + e^{\frac{(V-V_{0.5})}{k_i}}}$<br>$\tau^{\mathrm{f}}_{h_{\mathrm{SHL1}}}(V) = \tau_{h^{\mathrm{s}}_{\mathrm{SHL1}}}(V) = \frac{a}{1 + e^{\frac{(V-b)}{c}}} + d$<br>$I_{\mathrm{SHL1}} = \overline{g}_{\mathrm{SHL1}} \cdot m^3_{\mathrm{SHL1}} \cdot (0.7h^{\mathrm{f}}_{\mathrm{SHL1}} + 0.3h^{\mathrm{s}}_{\mathrm{SHL1}}) \cdot (V - V_K)$ |
| **KVS1** | $m_{\mathrm{KVS1},\infty}(V) = \frac{1}{1 + e^{\frac{-(V-V_{0.5})}{k_a}}}$<br>$h_{\mathrm{KVS1},\infty}(V) = \frac{1}{1 + e^{\frac{(V-V_{0.5})}{k_i}}}$<br>$\tau_{m_{\mathrm{KVS1}}}(V) = \tau_{h_{\mathrm{KVS1}}}(V) = \frac{a}{1 + e^{\frac{(V-b)}{c}}} + d$<br>$I_{\mathrm{KVS1}} = \overline{g}_{\mathrm{KVS1}} \cdot m_{\mathrm{KVS1}} \cdot h_{\mathrm{KVS1}} \cdot (V - V_K)$ |
| **SHK1** | $m_{\mathrm{SHK1},\infty}(V) = \frac{1}{1 + e^{\frac{-(V-V_{0.5})}{k_a}}}$<br>$\tau_{m_{\mathrm{SHK1}}}(V) = \frac{a}{e^{\frac{-(V-b)}{c}} + e^{\frac{(V-d)}{\tilde{e}}}} + f$<br>$h_{\mathrm{SHK1},\infty}(V) = \frac{1}{1 + e^{\frac{(V-V_{0.5})}{k_i}}}$<br>$\tau_{h_{\mathrm{SHK1}}} = a$<br>$I_{\mathrm{SHK1}} = \overline{g}_{SHK1} \cdot m_{\mathrm{SHK1}} \cdot h_{\mathrm{SHK1}} \cdot (V - V_K)$ |

| | |
|---|---|
| **KQT3** | $m^{\mathrm{f}}_{\mathrm{KQT3},\infty}(V) = m^{\mathrm{s}}_{\mathrm{KQT3},\infty}(V) = \frac{1}{1+e^{\frac{-(V-V_{0.5})}{k_a}}}$ <br> $\tau^{\mathrm{f}}_{m_{\mathrm{KQT3}}}(V) = \frac{a}{1+\left(\frac{V+b}{c}\right)^2}$ <br> $\tau^{\mathrm{s}}_{m_{\mathrm{KQT}}}(V) = a + \frac{b}{1+10^{-c(d-V)}} + \frac{\tilde{e}}{1+10^{-f(g+V)}}$ <br> $w_{\mathrm{KQT3},\infty}(V) = s_{\mathrm{KQT3},\infty}(V) = a + \frac{b}{1+e^{\frac{(V-V_{0.5})}{k_i}}}$ <br> $\tau_{w_{\mathrm{KQT3}}}(V) = a + \frac{b}{1+\left(\frac{V-c}{d}\right)^2}$ <br> $\tau_{s_{\mathrm{KQT3}}} = a$ <br> $I_{\mathrm{KQT3}} = \bar{g}_{\mathrm{KQT3}} \cdot \left(0.7m^{\mathrm{f}}_{\mathrm{KQT3}} + 0.3m^{\mathrm{s}}_{\mathrm{KQT3}}\right) \cdot w_{\mathrm{KQT3}} \cdot s_{\mathrm{KQT3}} \cdot (V - V_K)$ |
| **EGL2** | $m_{\mathrm{EGL2},\infty}(V) = \frac{1}{1+e^{\frac{-(V-V_{0.5})}{k_a}}}$ <br> $\tau_{m_{\mathrm{EGL2}}}(V) = \frac{a}{1+e^{\frac{(V-b)}{c}}} + d$ <br> $I_{\mathrm{EGL2}} = \bar{g}_{\mathrm{EGL2}} \cdot m_{\mathrm{EGL2}} \cdot (V - V_K)$ |
| **EGL36** | $m^{\mathrm{f}}_{\mathrm{EGL36},\infty}(V) = m^{\mathrm{m}}_{\mathrm{EGL36},\infty}(V) = m^{\mathrm{s}}_{\mathrm{EGL36},\infty}(V) = \frac{1}{1+e^{\frac{-(V-V_{0.5})}{k_a}}}$ <br> $\tau^{\mathrm{f}}_{m_{\mathrm{EGL36}}} = \tau^{\mathrm{m}}_{m_{\mathrm{EGL36}}} = \tau^{\mathrm{s}}_{m_{\mathrm{EGL36}}} = a$ <br> $I_{\mathrm{EGL36}} = \bar{g}_{\mathrm{EGL36}} \cdot \left(0.33m^{\mathrm{f}}_{\mathrm{EGL36}} + 0.36m^{\mathrm{m}}_{\mathrm{EGL36}} + 0.39m^{\mathrm{s}}_{\mathrm{EGL36}}\right) \cdot (V - V_K)$ |
| **IRK** | $m_{\mathrm{IRK},\infty}(V) = \frac{1}{1+e^{\frac{(V-V_{0.5})}{k_a}}}$ <br> $\tau_{m_{\mathrm{IRK}}}(V) = \frac{a}{e^{\frac{-(V-b)}{c}} + e^{\frac{(V-d)}{\tilde{e}}}} + f$ <br> $I_{\mathrm{IRK}} = \bar{g}_{\mathrm{IRK}} \cdot m_{\mathrm{IRK}} \cdot (V - V_K)$ |
| **EGL19** | $m_{EGL19,\infty}(V) = \frac{1}{1+e^{\frac{-(V-V_{0.5})}{k_a}}}$ <br> $\tau_{m_{EGL19}}(V) = \left[ae^{-\left(\frac{V-b}{c}\right)^2}\right] + \left[de^{-\left(\frac{V-\tilde{e}}{f}\right)^2}\right] + g$ <br> $h_{EGL19,\infty}(V) = \left[\frac{a}{1+e^{\frac{-(V-V_{0.5})}{k_i}}} + b\right] \cdot \left[\frac{c}{1+e^{\frac{(v-V^b_{0.5})}{k^b_i}}} + d\right]$ <br> $\tau_{h_{EGL19}}(V) = a\left[\frac{b}{1+e^{\frac{(V-c)}{d}}} + \frac{\tilde{e}}{1+e^{\frac{(V-f)}{g}}} + h\right]$ <br> $I_{EGL19} = \bar{g}_{EGL19} \cdot m_{EGL19} \cdot h_{EGL19} \cdot (V - V_{Ca})$ |

| | |
|---|---|
| **UNC2** | $m_{\mathrm{UNC2},\infty}(V) = \frac{1}{1+e^{\frac{-(V-V_{0.5})}{k_a}}}$<br>$\tau_{m_{\mathrm{UNC2}}}(V) = \frac{a}{e^{\frac{-(V-b)}{c}} + e^{\frac{(V-b)}{d}}} + \tilde{e}$<br>$h_{\mathrm{UNC2},\infty}(V) = \frac{1}{1+e^{\frac{(V-V_{0.5})}{k_i}}}$<br>$\tau_{h_{\mathrm{UNC2}}}(V) = \frac{a}{1+e^{\frac{-(V-b)}{c}}} + \frac{d}{1+e^{\frac{(V-\tilde{e})}{f}}}$<br>$I_{\mathrm{UNC2}} = \bar{g}_{\mathrm{UNC2}} \cdot m_{\mathrm{UNC2}} \cdot h_{\mathrm{UNC2}} \cdot (V - V_{Ca})$ |
| **CCA1** | $m_{\mathrm{CCA1},\infty}(V) = \frac{1}{1+e^{\frac{-(V-V_{0.5})}{k_a}}}$<br>$h_{\mathrm{CCA1},\infty}(V) = \frac{1}{1+e^{\frac{(V-V_{0.5})}{k_i}}}$<br>$\tau_{m_{\mathrm{CCA1}}}(V) = \frac{a}{1+e^{\frac{-(V-b)}{c}}} + d$<br>$\tau_{h_{\mathrm{CCA1}}}(V) = \frac{a}{1+e^{\frac{(V-b)}{c}}} + d$<br>$I_{\mathrm{CCA1}} = \bar{g}_{\mathrm{CCA1}} \cdot m_{\mathrm{CCA1}}^2 \cdot h_{\mathrm{CCA1}} \cdot (V - V_{Ca})$ |
| **SLO1_UNC2** | $m_{\mathrm{BK},\infty}(V, Ca) = \frac{m_{\mathrm{CaV}} k_o^+ (\alpha + \beta + k_c^-)}{(k_o^+ + k_o^-)(k_c^- + \alpha) + \beta k_c^-}$<br>$\tau_{m_{\mathrm{BK}}}(V, Ca) = \frac{\alpha + \beta + k_c^-}{(k_o^+ + k_o^-)(k_c^- + \alpha) + \beta k_c^-}$<br>$\alpha = \frac{m_{\mathrm{CaV},\infty}}{\tau_{m_{\mathrm{CaV}}}}$<br>$\beta = \tau_{m_{\mathrm{CaV}}}^{-1} - \alpha$<br>$I_{\mathrm{BK}} = \bar{g}_{\mathrm{BK}} \cdot m_{\mathrm{BK}} \cdot h_{\mathrm{CaV}} \cdot (V - V_K)$ |
| **SLO1_EGL19** | $m_{\mathrm{BK},\infty}(V, Ca) = \frac{m_{\mathrm{CaV}} k_o^+ (\alpha + \beta + k_c^-)}{(k_o^+ + k_o^-)(k_c^- + \alpha) + \beta k_c^-}$<br>$\tau_{m_{\mathrm{BK}}}(V, Ca) = \frac{\alpha + \beta + k_c^-}{(k_o^+ + k_o^-)(k_c^- + \alpha) + \beta k_c^-}$<br>$\alpha = \frac{m_{\mathrm{CaV},\infty}}{\tau_{m_{\mathrm{CaV}}}}$<br>$\beta = \tau_{m_{\mathrm{CaV}}}^{-1} - \alpha$<br>$I_{\mathrm{BK}} = \bar{g}_{\mathrm{BK}} \cdot m_{\mathrm{BK}} \cdot h_{\mathrm{CaV}} \cdot (V - V_K)$ |
| **SLO2_UNC2** | $m_{\mathrm{BK},\infty}(V, Ca) = \frac{m_{\mathrm{CaV}} k_o^+ (\alpha + \beta + k_c^-)}{(k_o^+ + k_o^-)(k_c^- + \alpha) + \beta k_c^-}$<br>$\tau_{m_{\mathrm{BK}}}(V, Ca) = \frac{\alpha + \beta + k_c^-}{(k_o^+ + k_o^-)(k_c^- + \alpha) + \beta k_c^-}$<br>$\alpha = \frac{m_{\mathrm{CaV},\infty}}{\tau_{m_{\mathrm{CaV}}}}$<br>$\beta = \tau_{m_{\mathrm{CaV}}}^{-1} - \alpha$<br>$I_{\mathrm{BK}} = \bar{g}_{\mathrm{BK}} \cdot m_{\mathrm{BK}} \cdot h_{\mathrm{CaV}} \cdot (V - V_K)$ |

| | |
|---|---|
| **SLO2_EGL19** | $m_{\mathrm{BK},\infty}(V, Ca) = \dfrac{m_{\mathrm{CaV}} k_o^+ (\alpha + \beta + k_c^-)}{(k_o^+ + k_o^-)(k_c^- + \alpha) + \beta k_c^-}$<br>$\tau_{m_{\mathrm{BK}}}(V, Ca) = \dfrac{\alpha + \beta + k_c^-}{(k_o^+ + k_o^-)(k_c^- + \alpha) + \beta k_c^-}$<br>$\alpha = \dfrac{m_{\mathrm{CaV},\infty}}{\tau_{m_{\mathrm{CaV}}}}$<br>$\beta = \tau_{m_{\mathrm{CaV}}}^{-1} - \alpha$<br>$I_{\mathrm{BK}} = \bar{g}_{\mathrm{BK}} \cdot m_{\mathrm{BK}} \cdot h_{\mathrm{CaV}} \cdot (V - V_K)$ |
| **KCNL** | $m_{\mathrm{KCNL},\infty}(Ca) = \dfrac{Ca}{K_{Ca} + Ca}$<br>$\tau_{m_{\mathrm{KCNL}}} = a$<br>$I_{\mathrm{KCNL}} = \bar{g}_{\mathrm{KCNL}} \cdot m_{\mathrm{KCNL}} \cdot (V - V_K)$ |
| **NCA** | $I_{NCA} = \bar{g}_{NCA} \cdot (V - V_{Na})$ |

**Table S2. Ion Channel Model Parameters**

| Ion Channel | Function | Parameter | Value | Unit |
|---|---|---|---|---|
| SHL1 | $m_\infty$ | $V_{0.5}$ | 11 | mV |
| | | $k_a$ | 14.1 | mV |
| | $h_\infty$ | $V_{0.5}$ | -33.1 | mV |
| | | $k_i$ | 8.3 | mV |
| | $\tau_m$ | a | 13.8 | ms |
| | | b | -17.5 | mV |
| | | c | 12.9 | mV |
| | | d | -3.7 | mV |
| | | e | 6.5 | mV |
| | | f | 1.9 | ms |
| | $\tau_h^f$ | a | 539.2 | ms |
| | | b | -28.2 | mV |
| | | c | 4.9 | mV |
| | | d | 27.3 | ms |
| | $\tau_h^s$ | a | 8422 | ms |
| | | b | -37.7 | mV |
| | | c | 6.4 | mV |
| | | d | 118.9 | ms |
| SHK1 | $m_\infty$ | $V_{0.5}$ | 20.4 | mV |
| | | $k_a$ | 7.7 | mV |
| | $h_\infty$ | $V_{0.5}$ | -7 | mV |
| | | $k_i$ | 5.8 | mV |
| | $\tau_m$ | a | 26.6 | ms |
| | | b | -33.7 | mV |
| | | c | 15.8 | mV |
| | | d | -33.7 | mV |
| | | e | 15.4 | mV |
| | | f | 2 | ms |
| | $\tau_h$ | a | 1400 | ms |
| KVS1 | $m_\infty$ | $V_{0.5}$ | 57.1 | mV |
| | | $k_a$ | 25 | mV |
| | $h_\infty$ | $V_{0.5}$ | 47 | mV |
| | | $k_i$ | 11.1 | mV |
| | $\tau_m$ | a | 30 | ms |
| | | b | 18.1 | mV |
| | | c | 20 | mV |
| | | d | 1 | ms |
| | $\tau_h^f$ | a | 88.5 | ms |
| | | b | 50 | mV |

| | | | | |
|---|---|---|---|---|
| | | c | -15 | mV |
| | | d | 53.4 | ms |
| **KQT3** | $m_\infty$ | $V_{0.5}$ | -12.6726 | mV |
| | | $k_a$ | 15.8008 | mV |
| | $w_\infty$ | $V_{0.5}$ | -1.084 | mV |
| | | $k_i$ | 28.78 | mV |
| | | a | 0.49 | |
| | | b | 0.51 | |
| | $s_\infty$ | $V_{0.5}$ | -45.3 | mV |
| | | $k_i$ | 12.3 | mV |
| | | a | 0.34 | |
| | | b | 0.66 | |
| | $\tau_m^f$ | a | 395.3 | ms |
| | | b | 38.1 | mV |
| | | c | 33.59 | mV |
| | $\tau_m^s$ | a | 5503 | ms |
| | | b | -5345.4 | ms |
| | | c | 0.02827 | $mV^{-1}$ |
| | | d | -23.9 | mV |
| | | e | 4590.6 | ms |
| | | f | 0.0357 | $mV^{-1}$ |
| | | g | 14.15 | mV |
| | $\tau_w$ | a | 0.544 | ms |
| | | b | 29.2 | ms |
| | | c | -48.09 | mV |
| | | d | 48.83 | mV |
| | $\tau_s$ | a | 500000 | ms |
| **EGL2** | $m_\infty$ | $V_{0.5}$ | -6.9 | mV |
| | | $k_a$ | 14.9 | mV |
| | $\tau_m$ | a | 1845.8 | ms |
| | | b | -122.6 | mV |
| | | c | -13.8 | mV |
| | | d | 1517.74 | ms |
| **EGL36** | $m_\infty$ | $V_{0.5}$ | 63 | mV |
| | | $k_a$ | 28.5 | mV |
| | $\tau_m^s$ | a | 355 | ms |
| | $\tau_m^m$ | a | 63 | ms |
| | $\tau_m^f$ | a | 13 | ms |
| **IRK** | $m_\infty$ | $V_{0.5}$ | -82 | mV |
| | | $k_a$ | 13 | mV |
| | $\tau_m$ | a | 17.1 | ms |
| | | b | -17.8 | mV |

| | | | | |
|---|---|---|---|---|
| | | c | 20.3 | mV |
| | | d | -43.4 | mV |
| | | e | 11.2 | mV |
| | | f | 3.8 | ms |
| **EGL19** | $m_\infty$ | $V_{0.5}$ | 5.6 | mV |
| | | $k_a$ | 7.5 | mV |
| | $h_\infty$ | $V_{0.5}$ | 24.9 | mV |
| | | $k_i$ | 12 | mV |
| | | $V_{0.5}{}^{b}$ | -20.5 | mV |
| | | $k_i{}^{b}$ | 8.1 | mV |
| | | a | 1.43 | |
| | | b | 0.14 | |
| | | c | 5.96 | |
| | | d | 0.6 | |
| | $\tau_m$ | a | 2.9 | ms |
| | | b | 5.2 | mV |
| | | c | 6 | mV |
| | | d | 1.9 | ms |
| | | e | 1.4 | mV |
| | | f | 30 | mV |
| | | g | 2.3 | ms |
| | $\tau_h$ | a | 0.4 | |
| | | b | 44.6 | ms |
| | | c | -23 | mV |
| | | d | 5 | mV |
| | | e | 36.4 | ms |
| | | f | 28.7 | mV |
| | | g | 3.7 | mV |
| | | h | 43.1 | ms |
| **UNC2** | $m_\infty$ | $V_{0.5}$ | -12.2 | mV |
| | | $k_a$ | 4 | mV |
| | $h_\infty$ | $V_{0.5}$ | -52.5 | mV |
| | | $k_i$ | 5.6 | mV |
| | $\tau_m$ | a | 1.5 | ms |
| | | b | -8.2 | mV |
| | | c | 9.1 | mV |
| | | d | 15.4 | mV |
| | | e | 0.1 | ms |
| | $\tau_h$ | a | 83.8 | ms |
| | | b | 52.9 | mV |
| | | c | -3.5 | mV |
| | | d | 72.1 | ms |

| | | | | |
|---|---|---|---|---|
| | | e | 23.9 | mV |
| | | f | -3.6 | mV |
| **CCA1** | $m_\infty$ | $V_{0.5}$ | -43.32 | mV |
| | | $k_a$ | 7.6 | mV |
| | $h_\infty$ | $V_{0.5}$ | -58 | mV |
| | | $k_i$ | 7 | mV |
| | $\tau_m$ | a | 40 | ms |
| | | b | -62.5 | mV |
| | | c | -12.6 | mV |
| | | d | 0.7 | ms |
| | $\tau_h$ | a | 280 | ms |
| | | b | -60.7 | mV |
| | | c | 8.5 | mV |
| | | d | 19.8 | ms |
| **SLO1** | $w_{yx}$ | | 0.013 | $mV^{-1}$ |
| | $w_{xy}$ | | -0.028 | $mV^{-1}$ |
| | $w_0^-$ | | 3.15 | $ms^{-1}$ |
| | $w_0^+$ | | 0.16 | $ms^{-1}$ |
| | $K_{xy}$ | | 55.73 | $\mu M/\mu m^3$ |
| | $n_{xy}$ | | 1.3 | |
| | $K_{yx}$ | | 34.34 | $\mu M/\mu m^3$ |
| | $n_{yx}$ | | 10−4 | |
| **SLO2** | $w_{yx}$ | | 0.019 | $mV^{-1}$ |
| | $w_{xy}$ | | -0.024 | $mV^{-1}$ |
| | $w_0^-$ | | 0.9 | $ms^{-1}$ |
| | $w_0^+$ | | 0.027 | $ms^{-1}$ |
| | $K_{xy}$ | | 93.45 | $\mu M/\mu m^3$ |
| | $n_{xy}$ | | 1.84 | |
| | $K_{yx}$ | | 3294.55 | $\mu M/\mu m^3$ |
| | $n_{yx}$ | | 10−5 | |
| **KCNL** | $K_{Ca}$ | | 0.33 | $\mu M$ |
| | $\tau_m$ | a | 6.3 | ms |
| **Intracellular calcium calculation** | $g_{sc}$ | | 40 | pS |
| | $V_{Ca}$ | | 60 | mV |
| | $r$ | | 13 | nm |
| | $F$ | | 96485 | $C\ mol^{-1}$ |
| | $D_{Ca}$ | | 250 | $\mu m^2/s$ |
| | $k_B^+$ | | 500 | $\mu M^{-1}\ s^{-1}$ |
| | $[B]_{tot}$ | | 30 | $\mu M$ |
| | $[Ca^{2+}]_{c,i}^n$ | | 0.05 | $\mu M$ |
| | $V_{cell}$ | | 31.63 | $\mu m^3$ |
| | $f$ | | 0.001 | |

| | | | |
|---|---|---|---|
| | $\tau_{Ca}$ | 50 | ms |
| | $[Ca^{2+}]_{eq}{}^{m}$ | 0.05 | $\mu$M/$\mu$m$^2$ |

**Table S3. Neuron Model Parameters**

| Parameters | AWC | AIY | RIM | VD5 | AVA |
|---|---|---|---|---|---|
| $R_a$ *(Ω·cm)* | 380 | 269 | 344 | 221 | 150 |
| $C_m$ *(μF/cm$^2$)* | 2 | 7 | 4 | 2 | 8 |
| $g_{pas}$ *(S/cm$^2$)* | 0.00003 | 0.000014 | 0.000077 | 0.00005 | 0.00007 |
| $R_m$ *(kΩ·cm$^2$)* | 33.3 | 71.4 | 13 | 20 | 14.3 |
| $e_{pas}$ *(mV)* | -65 | -54.5 | -33 | -75 | -33 |
| $\bar{g}_{SHL1}$ *(nS/μm$^2$)* | 0.0026 | 0.001 | 0.001 | 0.001 | 0.0004 |
| $\bar{g}_{SHK1}$ *(nS/μm$^2$)* | 0 | 0.00047 | 0.00015 | 0.0055 | 0.001 |
| $\bar{g}_{KVS1}$ *(nS/μm$^2$)* | 0 | 0.0005 | 0.0001 | 0 | 0 |
| $\bar{g}_{EGL2}$ *(nS/μm$^2$)* | 0 | 0.001 | 0.00001 | 0.001 | 0 |
| $\bar{g}_{EGL36}$ *(nS/μm$^2$)* | 0 | 0.00049 | 0.0012 | 0.0046 | 0 |
| $\bar{g}_{KQT3}$ *(nS/μm$^2$)* | 0.022 | 0.0005 | 0.00005 | 0.00016 | 0 |
| $\bar{g}_{EGL19}$ *(nS/μm$^2$)* | 0.006 | 0.0001 | 0 | 0.0001 | 0.002 |
| $\bar{g}_{UNC2}$ *(nS/μm$^2$)* | 0.001 | 0 | 0 | 0 | 0.0001 |
| $\bar{g}_{CCA1}$ *(nS/μm$^2$)* | 0 | 0.00017 | 0 | 0.0015 | 0.0006 |
| $\bar{g}_{SLO1\_EGL19}$ *(nS/μm$^2$))* | 0 | 0.00028 | 0.000013 | 0 | 0 |
| $\bar{g}_{SLO1\_UNC2}$ *(nS/μm$^2$)* | 0 | 0.000174 | 0.009 | 0.004 | 0.06 |
| $\bar{g}_{SLO2\_EGL19}$ *(nS/μm$^2$)* | 0 | 0.0015 | 0.00067 | 0.0013 | 0 |
| $\bar{g}_{SLO2\_UNC2}$ *(nS/μm$^2$)* | 0 | 0.0022 | 0 | 0.0006 | 0 |
| $\bar{g}_{KCNL}$*(nS/μm$^2$)* | 0 | 0.00063 | 0 | 0 | 0.005 |
| $\bar{g}_{NCA}$ *(nS/μm$^2$)* | 0 | 0.000085 | 0 | 0.00015 | 0.0007 |
| $\bar{g}_{IRK}$ *(nS/μm$^2$)* | 0 | 0.0011 | 0.00105 | 0 | 0 |

**Table S4. 302 Neurons and Functional Groups**

| Index | Neuron | Functional Group | Parameter Reference |
|---|---|---|---|
| 1 | I1L | interneuron | AIY |
| 2 | I1R | interneuron | AIY |
| 3 | I2L | interneuron | AIY |
| 4 | I2R | interneuron | AIY |
| 5 | I3 | interneuron | AIY |
| 6 | I4 | interneuron | AIY |
| 7 | I5 | interneuron | AIY |
| 8 | I6 | interneuron | AIY |
| 9 | M1 | body motor neuron | VD5 |
| 10 | M2L | body motor neuron | VD5 |
| 11 | M2R | body motor neuron | VD5 |
| 12 | M3L | body motor neuron | VD5 |
| 13 | M3R | body motor neuron | VD5 |
| 14 | M4 | body motor neuron | VD5 |
| 15 | M5 | body motor neuron | VD5 |
| 16 | MCL | body motor neuron | VD5 |
| 17 | MCR | body motor neuron | VD5 |
| 18 | MI | body motor neuron | VD5 |
| 19 | NSML | interneuron | AIY |
| 20 | NSMR | interneuron | AIY |
| 21 | ASIL | sensory neuron | AWC |
| 22 | ASIR | sensory neuron | AWC |
| 23 | ASJL | sensory neuron | AWC |
| 24 | ASJR | sensory neuron | AWC |
| 25 | AWAL | sensory neuron | AWC |
| 26 | AWAR | sensory neuron | AWC |
| 27 | ASGL | sensory neuron | AWC |
| 28 | ASGR | sensory neuron | AWC |
| 29 | AWBL | sensory neuron | AWC |
| 30 | AWBR | sensory neuron | AWC |
| 31 | ASEL | sensory neuron | AWC |
| 32 | ASER | sensory neuron | AWC |
| 33 | ADFL | sensory neuron | AWC |
| 34 | ADFR | sensory neuron | AWC |
| 35 | AFDL | sensory neuron | AWC |
| 36 | AFDR | sensory neuron | AWC |
| 37 | AWCL | sensory neuron | AWC |
| 38 | AWCR | sensory neuron | AWC |
| 39 | ASKL | sensory neuron | AWC |
| 40 | ASKR | sensory neuron | AWC |

| | | | |
|---|---|---|---|
| 41 | ASHL | sensory neuron | AWC |
| 42 | ASHR | sensory neuron | AWC |
| 43 | ADLL | sensory neuron | AWC |
| 44 | ADLR | sensory neuron | AWC |
| 45 | BAGL | sensory neuron | AWC |
| 46 | BAGR | sensory neuron | AWC |
| 47 | URXL | sensory neuron | AWC |
| 48 | URXR | sensory neuron | AWC |
| 49 | ALNL | sensory neuron | AWC |
| 50 | ALNR | sensory neuron | AWC |
| 51 | PLNL | sensory neuron | AWC |
| 52 | PLNR | sensory neuron | AWC |
| 53 | SDQL | sensory neuron | AWC |
| 54 | SDQR | sensory neuron | AWC |
| 55 | AQR | sensory neuron | AWC |
| 56 | PQR | sensory neuron | AWC |
| 57 | ALML | sensory neuron | AWC |
| 58 | ALMR | sensory neuron | AWC |
| 59 | AVM | sensory neuron | AWC |
| 60 | PVM | sensory neuron | AWC |
| 61 | PLML | sensory neuron | AWC |
| 62 | PLMR | sensory neuron | AWC |
| 63 | FLPL | sensory neuron | AWC |
| 64 | FLPR | sensory neuron | AWC |
| 65 | DVA | interneuron | AIY |
| 66 | PVDL | sensory neuron | AWC |
| 67 | PVDR | sensory neuron | AWC |
| 68 | ADEL | sensory neuron | AWC |
| 69 | ADER | sensory neuron | AWC |
| 70 | PDEL | sensory neuron | AWC |
| 71 | PDER | sensory neuron | AWC |
| 72 | PHAL | sensory neuron | AWC |
| 73 | PHAR | sensory neuron | AWC |
| 74 | PHBL | sensory neuron | AWC |
| 75 | PHBR | sensory neuron | AWC |
| 76 | PHCL | sensory neuron | AWC |
| 77 | PHCR | sensory neuron | AWC |
| 78 | IL2DL | sensory neuron | AWC |
| 79 | IL2DR | sensory neuron | AWC |
| 80 | IL2L | sensory neuron | AWC |
| 81 | IL2R | sensory neuron | AWC |
| 82 | IL2VL | sensory neuron | AWC |
| 83 | IL2VR | sensory neuron | AWC |

| | | | |
|---|---|---|---|
| 84 | CEPDL | sensory neuron | AWC |
| 85 | CEPDR | sensory neuron | AWC |
| 86 | CEPVL | sensory neuron | AWC |
| 87 | CEPVR | sensory neuron | AWC |
| 88 | URYDL | sensory neuron | AWC |
| 89 | URYDR | sensory neuron | AWC |
| 90 | URYVL | sensory neuron | AWC |
| 91 | URYVR | sensory neuron | AWC |
| 92 | OLLL | sensory neuron | AWC |
| 93 | OLLR | sensory neuron | AWC |
| 94 | OLQDL | sensory neuron | AWC |
| 95 | OLQDR | sensory neuron | AWC |
| 96 | OLQVL | sensory neuron | AWC |
| 97 | OLQVR | sensory neuron | AWC |
| 98 | IL1DL | sensory neuron | AWC |
| 99 | IL1DR | sensory neuron | AWC |
| 100 | IL1L | sensory neuron | AWC |
| 101 | IL1R | sensory neuron | AWC |
| 102 | IL1VL | sensory neuron | AWC |
| 103 | IL1VR | sensory neuron | AWC |
| 104 | AINL | interneuron | AIY |
| 105 | AINR | interneuron | AIY |
| 106 | AIML | interneuron | AIY |
| 107 | AIMR | interneuron | AIY |
| 108 | RIH | interneuron | AIY |
| 109 | URBL | interneuron | AIY |
| 110 | URBR | interneuron | AIY |
| 111 | RIR | interneuron | AIY |
| 112 | AIYL | interneuron | AIY |
| 113 | AIYR | interneuron | AIY |
| 114 | AIAL | interneuron | AIY |
| 115 | AIAR | interneuron | AIY |
| 116 | AUAL | interneuron | AIY |
| 117 | AUAR | interneuron | AIY |
| 118 | AIZL | interneuron | AIY |
| 119 | AIZR | interneuron | AIY |
| 120 | RIS | interneuron | AIY |
| 121 | ALA | interneuron | AIY |
| 122 | PVQL | interneuron | AIY |
| 123 | PVQR | interneuron | AIY |
| 124 | ADAL | interneuron | AIY |
| 125 | ADAR | interneuron | AIY |
| 126 | RIFL | interneuron | AIY |

| | | | |
|---|---|---|---|
| 127 | RIFR | interneuron | AIY |
| 128 | BDUL | interneuron | AIY |
| 129 | BDUR | interneuron | AIY |
| 130 | PVR | interneuron | AIY |
| 131 | AVFL | interneuron | AIY |
| 132 | AVFR | interneuron | AIY |
| 133 | AVHL | interneuron | AIY |
| 134 | AVHR | interneuron | AIY |
| 135 | PVPL | interneuron | AIY |
| 136 | PVPR | interneuron | AIY |
| 137 | LUAL | interneuron | AIY |
| 138 | LUAR | interneuron | AIY |
| 139 | PVNL | interneuron | AIY |
| 140 | PVNR | interneuron | AIY |
| 141 | AVG | interneuron | AIY |
| 142 | DVB | interneuron | AIY |
| 143 | RIBL | interneuron | AIY |
| 144 | RIBR | interneuron | AIY |
| 145 | RIGL | interneuron | AIY |
| 146 | RIGR | interneuron | AIY |
| 147 | RMGL | head motor neuron | RIM |
| 148 | RMGR | head motor neuron | RIM |
| 149 | AIBL | interneuron | AIY |
| 150 | AIBR | interneuron | AIY |
| 151 | RICL | interneuron | AIY |
| 152 | RICR | interneuron | AIY |
| 153 | SAADL | interneuron | AIY |
| 154 | SAADR | interneuron | AIY |
| 155 | SAAVL | interneuron | AIY |
| 156 | SAAVR | interneuron | AIY |
| 157 | AVKL | interneuron | AIY |
| 158 | AVKR | interneuron | AIY |
| 159 | DVC | interneuron | AIY |
| 160 | AVJL | interneuron | AIY |
| 161 | AVJR | interneuron | AIY |
| 162 | PVT | interneuron | AIY |
| 163 | AVDL | interneuron | AIY |
| 164 | AVDR | interneuron | AIY |
| 165 | AVL | interneuron | AIY |
| 166 | PVWL | interneuron | AIY |
| 167 | PVWR | interneuron | AIY |
| 168 | RIAL | interneuron | AIY |
| 169 | RIAR | interneuron | AIY |

| | | | |
|---|---|---|---|
| 170 | RIML | head motor neuron | RIM |
| 171 | RIMR | head motor neuron | RIM |
| 172 | AVEL | command neuron | AVA |
| 173 | AVER | command neuron | AVA |
| 174 | RMFL | head motor neuron | RIM |
| 175 | RMFR | head motor neuron | RIM |
| 176 | RID | interneuron | AIY |
| 177 | AVBL | command neuron | AVA |
| 178 | AVBR | command neuron | AVA |
| 179 | AVAL | command neuron | AVA |
| 180 | AVAR | command neuron | AVA |
| 181 | PVCL | command neuron | AVA |
| 182 | PVCR | command neuron | AVA |
| 183 | RIPL | interneuron | AIY |
| 184 | RIPR | interneuron | AIY |
| 185 | URADL | head motor neuron | RIM |
| 186 | URADR | head motor neuron | RIM |
| 187 | URAVL | head motor neuron | RIM |
| 188 | URAVR | head motor neuron | RIM |
| 189 | RMEL | head motor neuron | RIM |
| 190 | RMER | head motor neuron | RIM |
| 191 | RMED | head motor neuron | RIM |
| 192 | RMEV | head motor neuron | RIM |
| 193 | RMDDL | head motor neuron | RIM |
| 194 | RMDDR | head motor neuron | RIM |
| 195 | RMDL | head motor neuron | RIM |
| 196 | RMDR | head motor neuron | RIM |
| 197 | RMDVL | head motor neuron | RIM |
| 198 | RMDVR | head motor neuron | RIM |
| 199 | RIVL | head motor neuron | RIM |
| 200 | RIVR | head motor neuron | RIM |
| 201 | RMHL | head motor neuron | RIM |
| 202 | RMHR | head motor neuron | RIM |
| 203 | SABD | head motor neuron | RIM |
| 204 | SABVL | head motor neuron | RIM |
| 205 | SABVR | head motor neuron | RIM |
| 206 | SMDDL | head motor neuron | RIM |
| 207 | SMDDR | head motor neuron | RIM |
| 208 | SMDVL | head motor neuron | RIM |
| 209 | SMDVR | head motor neuron | RIM |
| 210 | SMBDL | head motor neuron | RIM |
| 211 | SMBDR | head motor neuron | RIM |
| 212 | SMBVL | head motor neuron | RIM |

| | | | |
|---|---|---|---|
| 213 | SMBVR | head motor neuron | RIM |
| 214 | SIBDL | head motor neuron | RIM |
| 215 | SIBDR | head motor neuron | RIM |
| 216 | SIBVL | head motor neuron | RIM |
| 217 | SIBVR | head motor neuron | RIM |
| 218 | SIADL | head motor neuron | RIM |
| 219 | SIADR | head motor neuron | RIM |
| 220 | SIAVL | head motor neuron | RIM |
| 221 | SIAVR | head motor neuron | RIM |
| 222 | DA1 | body motor neuron | VD5 |
| 223 | DA2 | body motor neuron | VD5 |
| 224 | DA3 | body motor neuron | VD5 |
| 225 | DA4 | body motor neuron | VD5 |
| 226 | DA5 | body motor neuron | VD5 |
| 227 | DA6 | body motor neuron | VD5 |
| 228 | DA7 | body motor neuron | VD5 |
| 229 | DA8 | body motor neuron | VD5 |
| 230 | DA9 | body motor neuron | VD5 |
| 231 | PDA | body motor neuron | VD5 |
| 232 | DB1 | body motor neuron | VD5 |
| 233 | DB2 | body motor neuron | VD5 |
| 234 | DB3 | body motor neuron | VD5 |
| 235 | DB4 | body motor neuron | VD5 |
| 236 | DB5 | body motor neuron | VD5 |
| 237 | DB6 | body motor neuron | VD5 |
| 238 | DB7 | body motor neuron | VD5 |
| 239 | AS1 | body motor neuron | VD5 |
| 240 | AS2 | body motor neuron | VD5 |
| 241 | AS3 | body motor neuron | VD5 |
| 242 | AS4 | body motor neuron | VD5 |
| 243 | AS5 | body motor neuron | VD5 |
| 244 | AS6 | body motor neuron | VD5 |
| 245 | AS7 | body motor neuron | VD5 |
| 246 | AS8 | body motor neuron | VD5 |
| 247 | AS9 | body motor neuron | VD5 |
| 248 | AS10 | body motor neuron | VD5 |
| 249 | AS11 | body motor neuron | VD5 |
| 250 | PDB | body motor neuron | VD5 |
| 251 | DD1 | body motor neuron | VD5 |
| 252 | DD2 | body motor neuron | VD5 |
| 253 | DD3 | body motor neuron | VD5 |
| 254 | DD4 | body motor neuron | VD5 |
| 255 | DD5 | body motor neuron | VD5 |

| | | | |
|---|---|---|---|
| 256 | DD6 | body motor neuron | VD5 |
| 257 | VA1 | body motor neuron | VD5 |
| 258 | VA2 | body motor neuron | VD5 |
| 259 | VA3 | body motor neuron | VD5 |
| 260 | VA4 | body motor neuron | VD5 |
| 261 | VA5 | body motor neuron | VD5 |
| 262 | VA6 | body motor neuron | VD5 |
| 263 | VA7 | body motor neuron | VD5 |
| 264 | VA8 | body motor neuron | VD5 |
| 265 | VA9 | body motor neuron | VD5 |
| 266 | VA10 | body motor neuron | VD5 |
| 267 | VA11 | body motor neuron | VD5 |
| 268 | VA12 | body motor neuron | VD5 |
| 269 | VB1 | body motor neuron | VD5 |
| 270 | VB2 | body motor neuron | VD5 |
| 271 | VB3 | body motor neuron | VD5 |
| 272 | VB4 | body motor neuron | VD5 |
| 273 | VB5 | body motor neuron | VD5 |
| 274 | VB6 | body motor neuron | VD5 |
| 275 | VB7 | body motor neuron | VD5 |
| 276 | VB8 | body motor neuron | VD5 |
| 277 | VB9 | body motor neuron | VD5 |
| 278 | VB10 | body motor neuron | VD5 |
| 279 | VB11 | body motor neuron | VD5 |
| 280 | VD1 | body motor neuron | VD5 |
| 281 | VD2 | body motor neuron | VD5 |
| 282 | VD3 | body motor neuron | VD5 |
| 283 | VD4 | body motor neuron | VD5 |
| 284 | VD5 | body motor neuron | VD5 |
| 285 | VD6 | body motor neuron | VD5 |
| 286 | VD7 | body motor neuron | VD5 |
| 287 | VD8 | body motor neuron | VD5 |
| 288 | VD9 | body motor neuron | VD5 |
| 289 | VD10 | body motor neuron | VD5 |
| 290 | VD11 | body motor neuron | VD5 |
| 291 | VD12 | body motor neuron | VD5 |
| 292 | VD13 | body motor neuron | VD5 |
| 293 | CANL | interneuron | AIY |
| 294 | CANR | interneuron | AIY |
| 295 | HSNL | body motor neuron | VD5 |
| 296 | HSNR | body motor neuron | VD5 |
| 297 | VC1 | body motor neuron | VD5 |
| 298 | VC2 | body motor neuron | VD5 |

| | | | |
|---|---|---|---|
| 299 | VC3 | body motor neuron | VD5 |
| 300 | VC4 | body motor neuron | VD5 |
| 301 | VC5 | body motor neuron | VD5 |
| 302 | VC6 | body motor neuron | VD5 |

**Table S5. Comparisons between Sibernetic and our body & environment model**

| | | OpenWorm Sibernetic | MetaWorm body & environment |
|---|---|---|---|
| Modelling | Worm body | ~34000-104000 particles | 984 vertices, 3341 tetrahedrons |
| | 3D scene scale (Worm body length) | 1.5-10 | ~1200 |
| Real-time Simulation | Simulation time per step (1 worm, 1 CPU core) | 0.4-2 s | 0.08-0.1 s |
| | Simulation time step (Default) | 0.00002 s | 1/240 s |
| Task & Behavior Analysis | Number of worms | One | One or more |
| | Ability for behavior analysis | No | Yes |

Siberntic is a physical simulator developed for simulations of *C. elegans* physical body dynamics within the OpenWorm Project[1]. Although Siberntic's particle model has advantages in some specific tasks, such as tasks related to pressure computation, MetaWorm's body & environment model outperforms it in many aspects, as demonstrated in Table S5. Firstly, from a modelling point of view, our tetrahedron worm body has much less elements compared to Sibernetic's particle body[2]. This results in a huge performance improvement while preserving anatomical authenticity. Moreover, with simplified hydrodynamics, scale of 3D simulation scene increased two orders compared to Sibernetic. Secondly, from a simulation perspective, the utilization of projective dynamics as the deformation solver has greatly reduced the simulation time per iteration step compared to Sibernetic. Moreover, projective dynamics exhibits stability even with large time steps, enabling the use of larger steps to accelerate simulation. Finally, by combining all the above advantages, we can perform task simulations of multiple worms or even worm populations, as shown in Movie S2. From the task point of view, this increased generality and applicability of tasks. Besides, the proposed TWRCS can numerically stably quantify worm locomotion, which is very useful for behavior analysis in worm tasks.

**Table S6. 136 Neurons Used in Simulation**

| No. | Index | Neuron | Recorded in Biological Experiment[3] | Network Input/Output |
|---|---|---|---|---|
| 1 | 39 | ASKL | yes | input |
| 2 | 40 | ASKR | yes | input |
| 3 | 49 | ALNL | yes | input |
| 4 | 50 | ALNR | yes | input |
| 5 | 61 | PLML | yes | input |
| 6 | 65 | DVA | yes | |
| 7 | 72 | PHAL | yes | input |
| 8 | 73 | PHAR | yes | input |
| 9 | 88 | URYDL | yes | input |
| 10 | 89 | URYDR | yes | input |
| 11 | 90 | URYVL | yes | input |
| 12 | 91 | URYVR | yes | input |
| 13 | 120 | RIS | yes | |
| 14 | 121 | ALA | yes | |
| 15 | 131 | AVFL | yes | |
| 16 | 132 | AVFR | yes | |
| 17 | 139 | PVNL | yes | |
| 18 | 140 | PVNR | yes | |
| 19 | 142 | DVB | yes | |
| 20 | 143 | RIBL | yes | |
| 21 | 144 | RIBR | yes | |
| 22 | 149 | AIBL | yes | |
| 23 | 150 | AIBR | yes | |
| 24 | 159 | DVC | yes | |
| 25 | 170 | RIML | yes | output |
| 26 | 171 | RIMR | yes | output |
| 27 | 172 | AVEL | yes | |
| 28 | 173 | AVER | yes | |
| 29 | 176 | RID | yes | |
| 30 | 177 | AVBL | yes | |
| 31 | 178 | AVBR | yes | |
| 32 | 179 | AVAL | yes | |
| 33 | 180 | AVAR | yes | |
| 34 | 189 | RMEL | yes | output |
| 35 | 190 | RMER | yes | output |
| 36 | 191 | RMED | yes | output |
| 37 | 192 | RMEV | yes | output |
| 38 | 199 | RIVL | yes | output |
| 39 | 200 | RIVR | yes | output |

| | | | | |
|---|---|---|---|---|
| 40 | 203 | SABD | yes | |
| 41 | 204 | SABVL | yes | |
| 42 | 205 | SABVR | yes | |
| 43 | 206 | SMDDL | yes | output |
| 44 | 207 | SMDDR | yes | output |
| 45 | 208 | SMDVL | yes | output |
| 46 | 209 | SMDVR | yes | output |
| 47 | 218 | SIADL | yes | |
| 48 | 219 | SIADR | yes | |
| 49 | 220 | SIAVL | yes | |
| 50 | 221 | SIAVR | yes | |
| 51 | 222 | DA1 | yes | output |
| 52 | 228 | DA7 | yes | output |
| 53 | 230 | DA9 | yes | output |
| 54 | 231 | PDA | yes | |
| 55 | 232 | DB1 | yes | output |
| 56 | 233 | DB2 | yes | output |
| 57 | 238 | DB7 | yes | output |
| 58 | 248 | AS10 | yes | |
| 59 | 257 | VA1 | yes | output |
| 60 | 267 | VA11 | yes | output |
| 61 | 268 | VA12 | yes | output |
| 62 | 269 | VB1 | yes | output |
| 63 | 270 | VB2 | yes | output |
| 64 | 279 | VB11 | yes | output |
| 65 | 292 | VD13 | yes | output |
| 66 | 25 | AWAL | no | input |
| 67 | 26 | AWAR | no | input |
| 68 | 37 | AWCL | no | input |
| 69 | 38 | AWCR | no | input |
| 70 | 112 | AIYL | no | |
| 71 | 113 | AIYR | no | |
| 72 | 114 | AIAL | no | |
| 73 | 115 | AIAR | no | |
| 74 | 118 | AIZL | no | |
| 75 | 119 | AIZR | no | |
| 76 | 153 | SAADL | no | |
| 77 | 154 | SAADR | no | |
| 78 | 155 | SAAVL | no | |
| 79 | 156 | SAAVR | no | |
| 80 | 181 | PVCL | no | |
| 81 | 182 | PVCR | no | |
| 82 | 193 | RMDDL | no | output |

| | | | | |
|---|---|---|---|---|
| 83 | 194 | RMDDR | no | output |
| 84 | 195 | RMDL | no | output |
| 85 | 196 | RMDR | no | output |
| 86 | 197 | RMDVL | no | output |
| 87 | 198 | RMDVR | no | output |
| 88 | 210 | SMBDL | no | output |
| 89 | 211 | SMBDR | no | output |
| 90 | 212 | SMBVL | no | output |
| 91 | 213 | SMBVR | no | output |
| 92 | 223 | DA2 | no | output |
| 93 | 224 | DA3 | no | output |
| 94 | 225 | DA4 | no | output |
| 95 | 226 | DA5 | no | output |
| 96 | 227 | DA6 | no | output |
| 97 | 229 | DA8 | no | output |
| 98 | 234 | DB3 | no | output |
| 99 | 235 | DB4 | no | output |
| 100 | 236 | DB5 | no | output |
| 101 | 237 | DB6 | no | output |
| 102 | 251 | DD1 | no | output |
| 103 | 252 | DD2 | no | output |
| 104 | 253 | DD3 | no | output |
| 105 | 254 | DD4 | no | output |
| 106 | 255 | DD5 | no | output |
| 107 | 256 | DD6 | no | output |
| 108 | 258 | VA2 | no | output |
| 109 | 259 | VA3 | no | output |
| 110 | 260 | VA4 | no | output |
| 111 | 261 | VA5 | no | output |
| 112 | 262 | VA6 | no | output |
| 113 | 263 | VA7 | no | output |
| 114 | 264 | VA8 | no | output |
| 115 | 265 | VA9 | no | output |
| 116 | 266 | VA10 | no | output |
| 117 | 271 | VB3 | no | output |
| 118 | 272 | VB4 | no | output |
| 119 | 273 | VB5 | no | output |
| 120 | 274 | VB6 | no | output |
| 121 | 275 | VB7 | no | output |
| 122 | 276 | VB8 | no | output |
| 123 | 277 | VB9 | no | output |
| 124 | 278 | VB10 | no | output |
| 125 | 280 | VD1 | no | output |

| | | | | |
|---|---|---|---|---|
| 126 | 281 | VD2 | no | output |
| 127 | 282 | VD3 | no | output |
| 128 | 283 | VD4 | no | output |
| 129 | 284 | VD5 | no | output |
| 130 | 285 | VD6 | no | output |
| 131 | 286 | VD7 | no | output |
| 132 | 287 | VD8 | no | output |
| 133 | 288 | VD9 | no | output |
| 134 | 289 | VD10 | no | output |
| 135 | 290 | VD11 | no | output |
| 136 | 291 | VD12 | no | output |

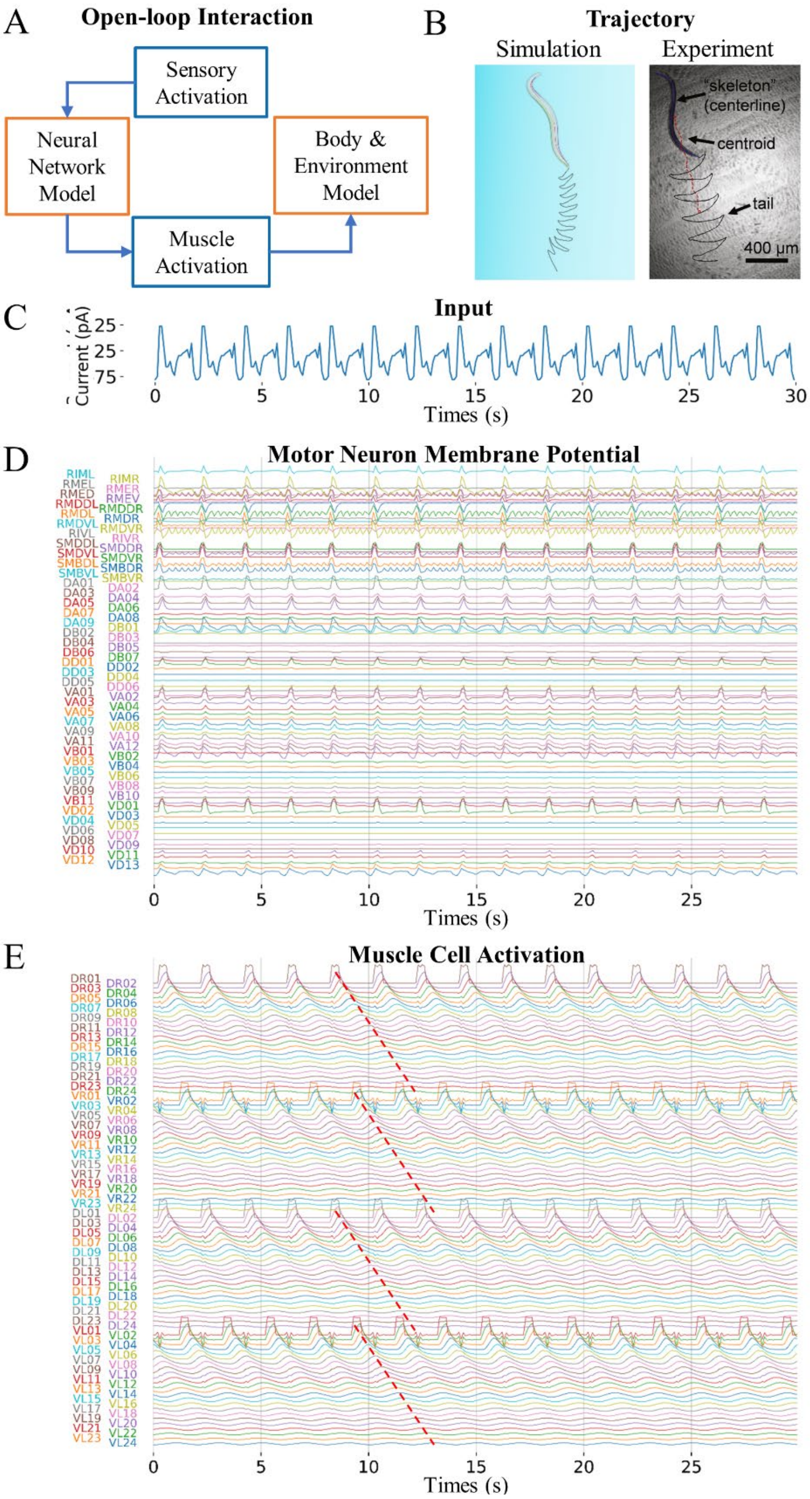


**Figure S1. The open-loop 3D simulation of C. elegans locomotion behavior.**

**A.** The information flow diagram of the open-loop interaction between neural network model and body & environment model. **B.** Comparison of *C. elegans* trajectory during forward locomotion in simulation (left) and experiment (right, from a published article[4]). **C.** Input current injected to sensory neurons during locomotion. **D.** Left, neural network model of *C. elegans*. Right, membrane potentials of all motor neurons in the model during locomotion. **E.** Left, the body model with 96 muscle cells. Right, the activation signals of all muscle cells during locomotion. The dashed red line indicates that the activation signal shifting from head to tail.

**Movie S1.** Closed-loop 3D simulation of *C. elegans* locomotion behavior.

**Movie S2.** Closed-loop 3D simulation of six *C. elegans* locomotion behavior.

**Movie S3.** Open-loop 3D simulation of *C. elegans* locomotion behavior.
